# Overnight fasting facilitates safety learning by changing the neurophysiological response to relief from threat omission

**DOI:** 10.1101/2023.09.06.556396

**Authors:** Silvia Papalini, Tom Beckers, Lukas Van Oudenhove, Bram Vervliet

**Author notes:** Corresponding author, Email: Dr., Tiensestraat 102 Leuven (3000), Belgium.

## Abstract

Excessive avoidance and slow extinction of fear are hallmarks of anxiety disorders. We have previously found that overnight fasting diminishes excessive avoidance and speeds up fear extinction by decreasing subjective relief during threat omissions. Since relief tracks the reward prediction error signal that governs safety learning, we hypothesized that these effects of fasting might be linked to a decreased activation in brain regions related to reward prediction error processing. Hence, we replicated our previous study in a 3T-MRI scanner. Overnight fasting increased effective avoidance and sped up fear extinction learning. During extinction, the fasting group showed lower activations in the ventromedial prefrontal cortex and nucleus accumbens in response to threat omissions signaled by a safe cue. Nucleus accumbens activations were modulated by relief in the control group. This study provides support for overnight fasting as an adjunct to treatments for anxiety, but the effects should be investigated in anxious patients.

## Introduction

The ability to identify and avoid cues that predict threat is essential to behavioral adaptation and, ultimately, survival. However, when former threat cues no longer predict threat, fear and avoidance should also decrease accordingly. In clients with anxiety disorders, a top leading cause of mental burden worldwide^1^, fear and avoidance tend to persist even in situations of objective safety. The standard treatment for these disorders, exposure therapy, decreases excessive fear and avoidance by gradually approaching these situations and promoting safety learning. Although this therapy is considered effective, nearly half of these clients do not benefit from this intervention^2^. Therefore, gaining insight in behavioral and neural mechanisms of safety learning, as well as discovering new ways to enhance safety learning, is of great clinical relevance.

Fear extinction studies in rodents have found that acute fasting enhances safety learning^3,4^. In the same vein, we recently observed in healthy humans that overnight fasting optimizes two forms of safety learning: active avoidance learning and fear extinction learning. Specifically, we found that overnight fasting reduced unnecessary avoidance during a signal of safety, while maintaining effective avoidance during a signal of upcoming threat. This decrease in unnecessary avoidance was mediated by lower levels of relief pleasantness reported by fasted versus fed participants during omissions of threat signaled by a safe cue. Interestingly, we also observed a faster decline in relief pleasantness ratings during fear extinction, suggesting faster decrease of threat expectancies^5^. Subjective relief pleasantness is the emotional experience of a better-than-expected outcome in the context of threat omission^6–8^ and has therefore been related to the reward prediction error, rPE^9–11^, which is assumed to reinforce avoidance actions^8^. Thus, it seems that overnight fasting decreased the reward value of safety (less relief pleasantness) and decreased the motivation to achieve safety (less avoidance)^5^. Together with the obvious fact that fasting increases the value of food as well as the motivation to achieve food^12–14^, these results may indicate fasting biases an animal or human to focus on food at the expense of safety. This trade-off may be adaptive for successful foraging: searching for food implies leaving the safe nest.

The current functional Magnetic Resonance Imaging (fMRI) study was set up to examine whether the decreased relief ratings in avoidance learning and fear extinction are indeed indicative of decreased reward activations during omission of threat. Rodent studies have previously shown that unexpected, but not expected, omissions of threat trigger dopaminergic activations in the Ventral Tegmental Area (VTA), similar to a rPE signal^15,16^. Via mesolimbic dopamine projections, these VTA activations probably recruit the nuclei accumbens (Nac), amygdala and hippocampus in concert to induce the valence-related plastic changes necessary for the formation of safety memories in fear extinction^17^ and avoidance learning^18,19^. Accordingly, human fMRI studies^20,21^ have found (patterns of) activations in the ventral striatum and the VTA during threat omissions that might contribute to establishing a new safety CS–>noUS memory that reduces the initial fear responses^22^. Hence, we hypothesized, based on our previous relief findings, that overnight fasting would decrease activation in these brain areas during threat omissions in avoidance learning and fear extinction. Because overnight fasting decreased relief ratings particularly during omissions after safety signals and unnecessary avoidance actions, we focused on these events to test our central hypothesis (as preregistered on the Open Science Framework, osf.io/yx8bq).

As in our previous experiment, we asked a Re-feeding Group (breakfast after overnight fasting) and a Fasting Group (14 to 16 hours of overnight fasting) to perform a short version of the Avoidance Relief Task, ART^5^, see **figure 1A**. This task starts with a Pavlovian fear conditioning phase, in which different visual stimuli (conditional stimuli, CSs) are paired (CS+) or not (CS-) with an unpleasant electrical stimulation (unconditioned stimulus, US). During the avoidance conditioning phase, participants can learn to press a button to avoid the unpleasant US. Finally, during the fear extinction and response prevention phase, the button is removed and the CSs are presented in the absence of the US. In all phases, subjective ratings of relief pleasantness, skin conductance reactivity (SCR), and brain activations (blood-oxygen-level-dependent, BOLD, signal) are measured during US omissions at CS offsets, to track correlates of the rPE signal that drives safety learning during differential conditioning, avoidance learning and fear extinction, see **figure 1B**.

**Figure 1.**
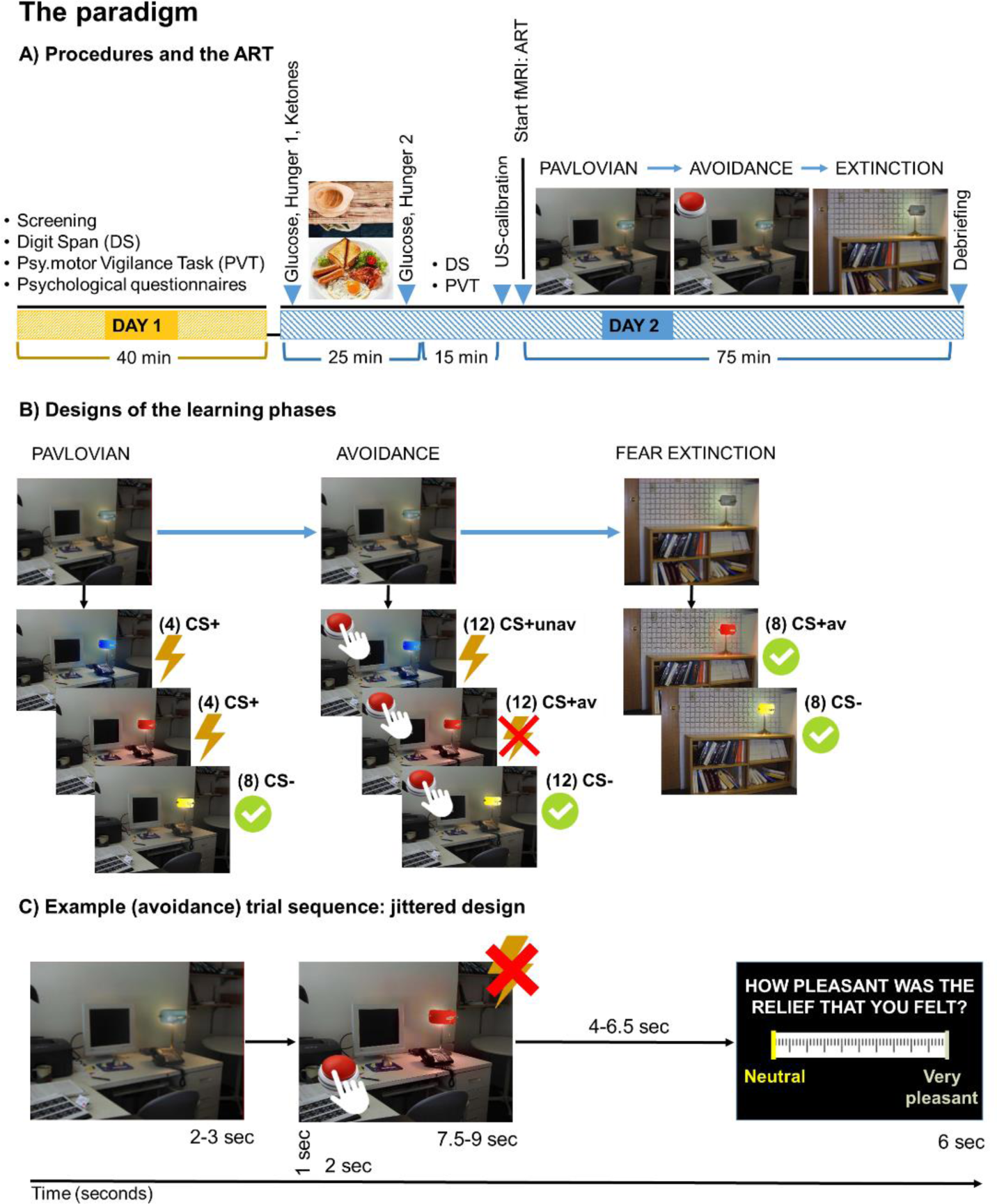
Overview of the procedures of the fMRI study and of the ART. **A)** During the first (day 1) and second (day 2) visits, we measured working memory capacity, psychomotor visual attention, and cognitive effort levels through the Digit Span (DS)^23^, the Psychomotor Visual Task (PVT)^24^, and the Need for Cognition Scale (NFS)^25^. These measurements were performed to exclude fasting effects on learning following changes in general cognitive skills. During the second visit, glucose levels were measured before and after breakfast in the Re-feeding Group and with a time interval of 20/25 minutes in the Fasting Group. Ketone levels were measured once at the beginning of the second session. After the neuropsychological assessment, participants were asked to lie inside the scanner. The intensity of the US was then calibrated following standardized procedures ^26^. The US was set to an intensity for which participants refer to the electrical stimulation as highly unpleasant but not painful. Following explanations about the task and relief ratings, the fMRI exam started for a total duration of about 75 minutes. After the acquisition of the anatomical scans, the participants performed the ART. Since we focused on the dynamics of learning, in the present fMRI study we did not include the test phases of the ART, as in ^5^. After fMRI acquisition, an explanation of hypotheses and aims was provided to each participant. **B)** The ART consists of three learning phases, a Pavlovian phase that uses a block-design, an avoidance learning and fear extinction learning phases that use an event-related type of presentation of the stimuli, see ^5^ for details. **C)** We applied a jittered design to better distinguish BOLD and SCR responses during US omissions. The illustration shows an example of a trial during the avoidance learning phase, where background, CS presentation (US anticipation), and CS offset (relief) were jittered events before that the online relief rating was collected.

We previously found that overnight fasting reduces avoidance and relief mostly to a safe CS-^5^. Therefore, we focused our hypotheses on fasting-induced changes to rPE-like processing of US omissions following CS-presentations. For the **avoidance conditioning phase**, we hypothesized that overnight fasting decreases Nac, VTA, and ventro medial Prefrontal Cortex (vmPFC) activations during US omissions following a CS-, as compared to US omissions following an actively avoided CS+ (Hypothesis 1-a). We further hypothesized that these weaker brain activations in the fasting group would be related to lower levels of avoidance during CS-, as compared to CS+ (Hypothesis 1-b), and also related to a reduction in subjective relief ratings (Hypothesis 1-c). For the **fear extinction phase**, we hypothesized that overnight fasting would decrease Nac, VTA, and vmPFC activations during US omissions following the CS-, as compared to US omissions following the CS+ (Hypothesis 2-a). We further hypothesized that these weaker brain activations would be related to a faster reduction in relief pleasantness ratings for CS-versus CS+ (Hypothesized 2-b). These a priori hypotheses (and further explorative tests) were preregistered at osf.io/yx8bq.

## Results

In this section, we only reported the principal contrasts of interest to test the pre-registered hypotheses, which include the Group by CS interactions from the (Generalized) Linea Mixed Models (GLMM, LMM) and two sample t-tests. We also included eventual main effects of Group, or Group by Trial interactions that were not pre-registered. The factor Trial was de-meaned to focus on the learning curve. Importantly, differently from the behavioral data and in line with the recent neuroimaging literature on rPE during threat omission^20,21^, we did not find the expected differentiation between brain activations to US omissions signaled by the CS+av versus CS-in either the extinction or the avoidance learning phase (in any of our Regions Of Interests, ROIs) unless group effects were taken into account. In the supplementary material, we included the results from the Pavlovian, avoidance, and fear extinction learning phases of the ART that involve ratings, brain activations, and SCRs during US anticipation/expectancy. These results were in line with the results from previous fear conditioning studies^20,27,28^. No group differences were found during anticipation.

We also investigated for group differences in explorative time-related analysis on the ROIs. In one type of time-related analysis, we first run a t-test on early compared to late omissions (CS-versus CS+av)early versus (CS-versus CS+av)late. For a second type of analysis, which can be intended as a growth curve modeling, we run a LMM for ROI including ‘Trial’ as covariate of interest. The results from these analyses replicated what already found in the main analysis without any significant group by time-related effect. Hence, these results are not reported here.

The expected effects of the fasting and re-feeding procedure on glucose and hunger levels as well as the results from the psychometric questionnaires (which -successfully-served to exclude effects of fasting on cognition) are also reported in the supplementary material. No analysis of the ketone levels was performed, since we did not detect any increase after overnight fasting (all values equal to zero).

### Avoidance

#### Ratings

*Relief-pleasantness ratings:* the results from the Group (FG/RG) by CS (CS-/CS+av) by Trial LMM analysis revealed the unexpected absence of a significant main effect of Group or interaction with Group (all *p-*values > 0.05), **figure 2B**, right panel.

**Figure 2.**
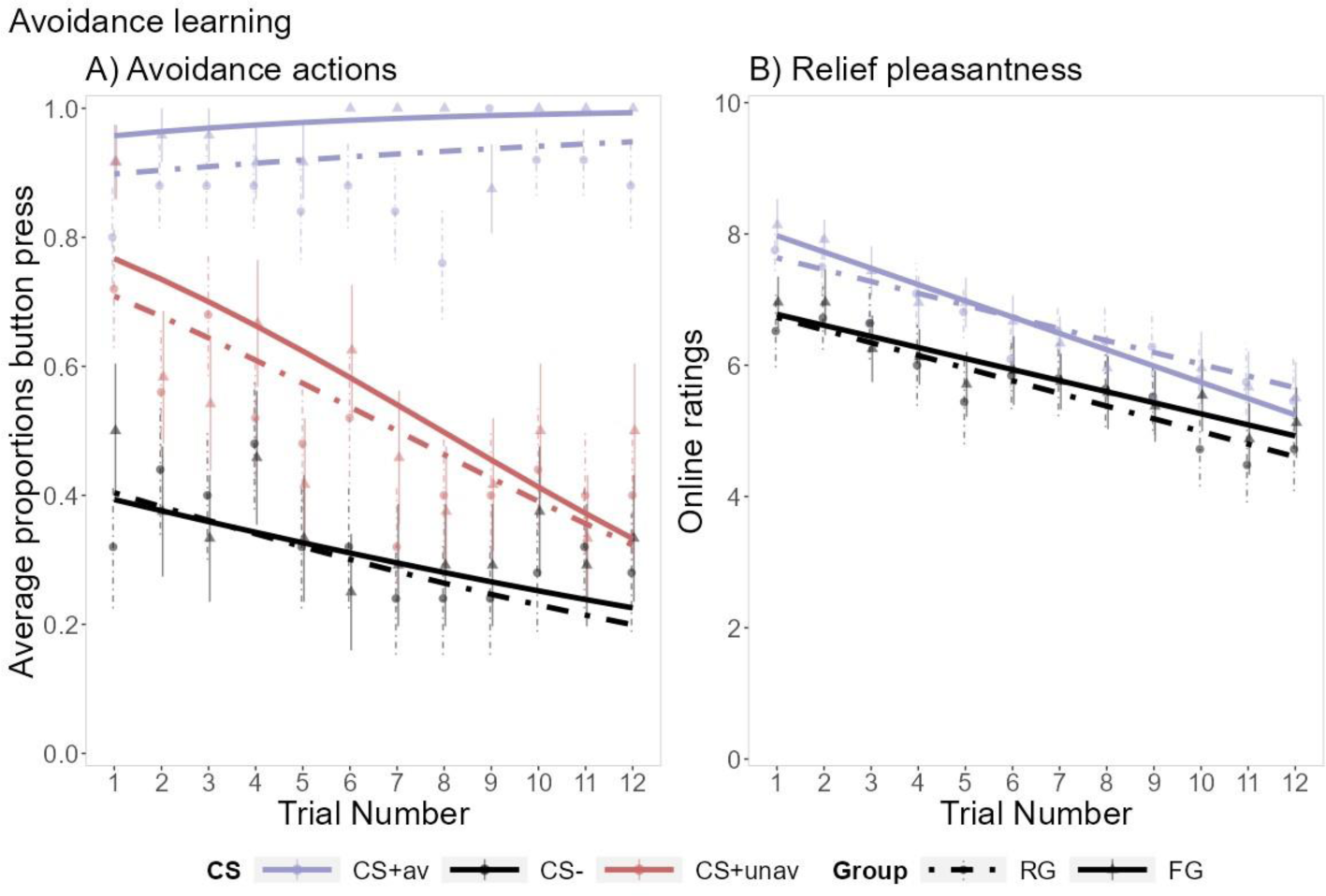
**A)** Avoidance actions and **B)** Relief pleasantness ratings during the avoidance phase for the Fasting Group (FG, continuous lines) and the Re-feeding Group (RG, dashed lines). **A)** The results from the GLMM analysis indicated a (general) lower probability of pressing the button for the CS- and CS+_unav_ (all *p*-values < 0.001), and a significant reduction in the probability to press the button over the trials of the CS- and CS+_unav_ when compared to the CS+_av_ (all p-values < 0.024). The Fasting Group, compared to the Re-feeding Group, showed a significantly higher probability to avoid the CS_+av_. **B)** Individual relief ratings were higher at the offset of the CS_+av_ and followed a rPE-like learning trajectory: high relief ratings during the first unexpected omissions of the US and lower ratings recorded at the end of the learning phase (Trial: *β* = - 0.180, ± 0.055, df = 48.141, t = - 3.287, *p* = 0.002, %95 CIs [- 0.287:- 0.072], RESI = 0.386). There was also an expected significantly lower relief for the safe CS- versus the CS+_av_ (CS: *β* = - 0.965, ± 0.287, df = 47.954, t = - 3.367, *p* = 0.002, %95 CIs [- 1.526:- 0.404], RESI = 0.425). No Group-related differences were found in the relief ratings recorded for each CS. This model also included the avoidance actions performed for the CS-. The results did not change when this covariate was removed from the analysis. Error bars represent the standard error of the mean, while the lines represent the models fit.

#### Behavioral data

*Avoidance actions*: results from the Group (FG/RG) by CS (CS- /CS+av/CS+unav) by Trial GLMM analysis revealed the expected significant Group by CS interaction (omnibus: *ꭓ^2^*= 8.955, df = 2, *p* = 0.011). The results from the planned contrasts indicated that compared to the Re-feeding group, the Fasting Group showed a lower probability to avoid the CS-than the CS+av (*β* = - 1.370, ± 0.458, z = - 2.995, *p* = 0.003, %95 CIs [- 2.303:- 0.503], Exp-*β* = 1.877) and a lower probability to avoid the CS+unav than the CS+av (*β* = - 1.211, ± 0.439, z = - 2.758, *p* = .006, %95 CIs [- 2.108:- 0.382], Exp-*β* = 1.467). We also found an unexpected higher probability to avoid the CS+av (which button presses for this CS were used as baseline) in the Fasting Group compared to the Re-feeding Group (*β* = 1.108, ± 0.578, z = 2.436, *p* = 0.015, %95 CIs [0.285:2.573], Exp-*β* = 1.228). This means that the expected Group by CS interaction reported above was dependent on an increased probability to effectively avoid the CS+av in the Fasting Group compared to Re-feeding group (and not by a decreased probability in unnecessary and ineffective avoidance in the Fasting group as we found in our previous study), see left panel in **figure 2A** for models fit.

Since we observed a large variance for the avoidance actions during the CS-, we additionally ran the same GLMM analysis taking into account the ‘better safe than sorry’ strategy. To do this, we calculated the individual frequency to press the button for the CS-. When we added this continuous covariate to the original GLMM the results did not change, confirming that the Fasting Group, independent of individual strategy, avoided in response to the CS+av more often than the Re- feeding Group.

Importantly, the result of an explorative GLMM analysis (Group: FG/RG by CS: CS-/CS+av/CS+unav, by Study design: fMRI/behavioral) indicated that the present Fasting Group showed a higher general probability to avoid than the Fasting Group examined in the previous behavioral study (*β* = -1.644, ± 0.492, z = -3.344, *p* < 0.001, 95% CIs [- 2.655:- 0.713], Exp-*β* = 2.703), **figure 3** for models fit.

**Figure 3.**
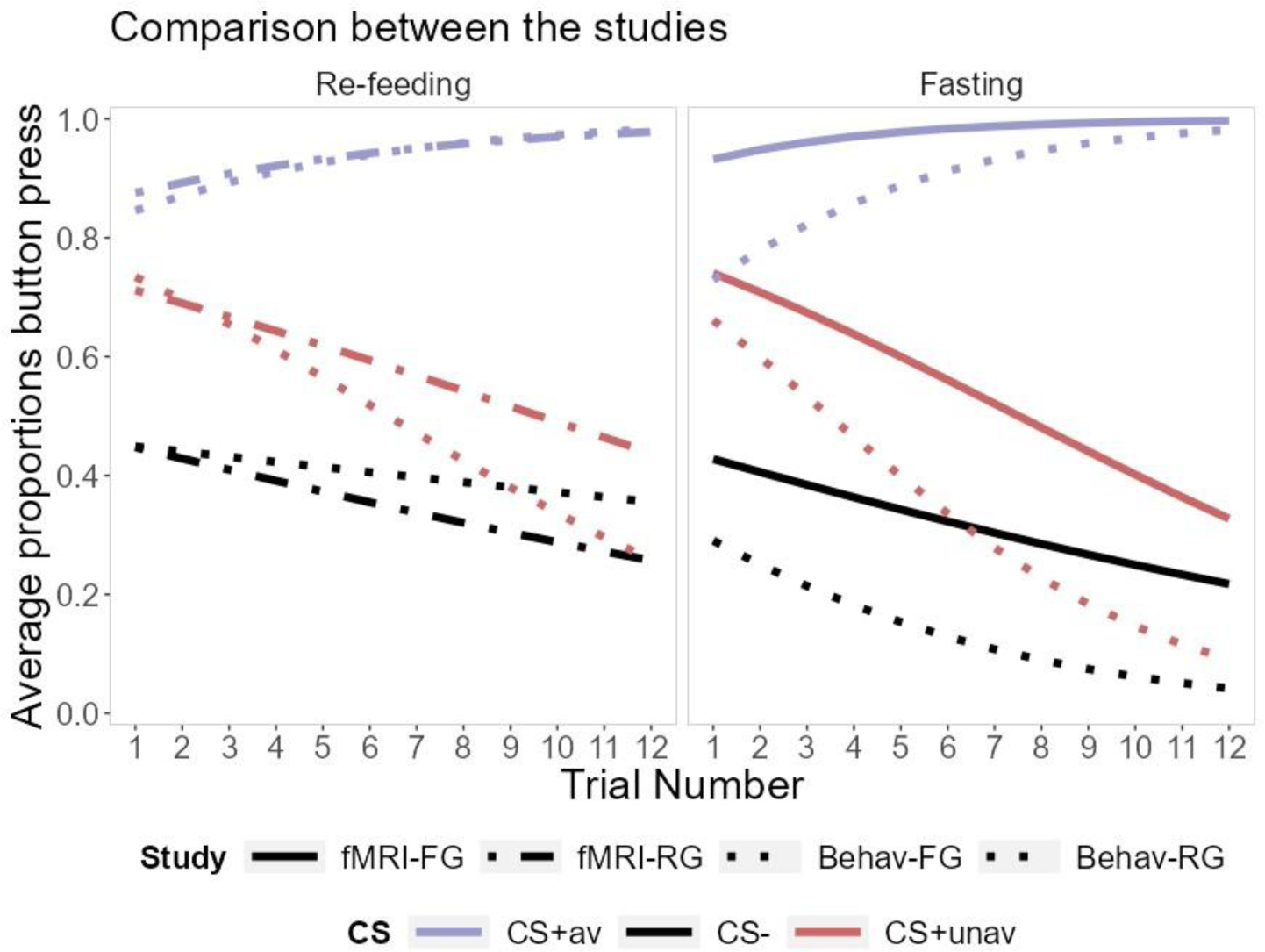
The avoidance profiles of the Fasting Group *(left panel)* and the Re-feeding Group *(right panel)*, during the previous behavioral (dashed lines) and the present fMRI (continuous lines) studies. The Fasting Group showed an increase in general avoidance behaviors in the MRI environment.

#### Neural data

##### A priori hypothesis

###### Group differences during US omission (ROIs)

In our original hypothesis **(hypothesis 1-a)**, we expected the Fasting Group to show a decreased VTA, Nac, and vmPFC activation in response to US omissions (CS-versus CS+av offset).

The result from a two sample t-test showed a trend towards a significant expected effect in the Nac (CS- vs CS+av: t = - 1.653, df = 39.764, *p* = 0.053, 95% CIs [- Inf:0.001], *d* = 0.498), which was driven by a lower brain activity in response to CS- offset in the Fasting Group compared to the Re-feeding Group (CS-: t = - 2.502, df = 41.675, *p* = 0.016, %95 CI s [- 0.153:- 0.016], *d* = 0.754). No other significant effects were found in the other two ROIs.

Given the absence of a group effect at the brain level there was no reason to test our **Hypothesis 1-b.** Similarly, given the lack of a significant group difference in relief levels during US omission, there was no reason for any mediation analyses **(hypothesis 1-c).**

#### Skin Conductance Responses

In a secondary analysis, we investigated if fasting changes SCR during US omission. The results from the GLMM (logistic component) showed no significant results (all *p*-values > 0.05). We followed up with a LMM (normal component with normalized non-zero values of the SCR) and found a significant main effect of Group (omnibus: F = 5.707, df = 1, *p* = 0.022). To characterize this Group effect, since we did not plan contrasts for a general main effect of Group, we followed up with a post hoc LMM analysis run separately for each CS type. The results indicated that compared to the Re-feeding Group, the Fasting Group showed a lower SCR to US omissions following both the CS- (*β* = - 0.351, ± 0.166, df = 28.225, t = - 2.111, *p* = 0.044, 95% CIs [- 0.668:- 0.019], RESI = 0.302) and the CS+av (*β* = - 0.342, ± 0.166, df = 34.976, t = - 2.061, *p* = 0.047, 95% CIs [- 0.684:- 0.090], RESI = 0.291), which, however, did not survive to Bonferroni correction.

### Extinction

Our second hypothesis was that overnight fasting would decrease VTA, Nac, and vmPFC responses to the CS- compared to the CS+av (**hypothesis 2-a**).

#### Ratings

*Relief pleasantness* ratings: as expected, the Group (FG/RG) by CS(CS-/CS+av) by Trial LMM analysis showed a significant Group by Trial interaction (omnibus: F = 8.903, df = 1, *p* = 0.005) and a Group by CS interaction (omnibus: F = 7.757, df = 1, *p* = 0.006). To further characterize these significant interactions, since we did not plan contrasts for an effect of Trial, we followed up with a post hoc LMM analysis run separately for each CS type. The results showed that compared to the Re-feeding group, relief ratings decreasing significantly faster in the Fasting Group for both the CS- (*β* = - 0.327, ± 0.009, t = - 3.305, df = 43.992, *p* = 0.001, 95% CIs [- 0.521:- 0.133], RESI= 0.453) and the CS+av (*β* = - 0.268, ± 0.115, t = - 2.334, df = 43.999, *p* = 0.024, 95% CIs [- 0.492:- 0.043], RESI = 0.303) in absence of any main effect of Group, see **figure 4**.

**Figure 4.**
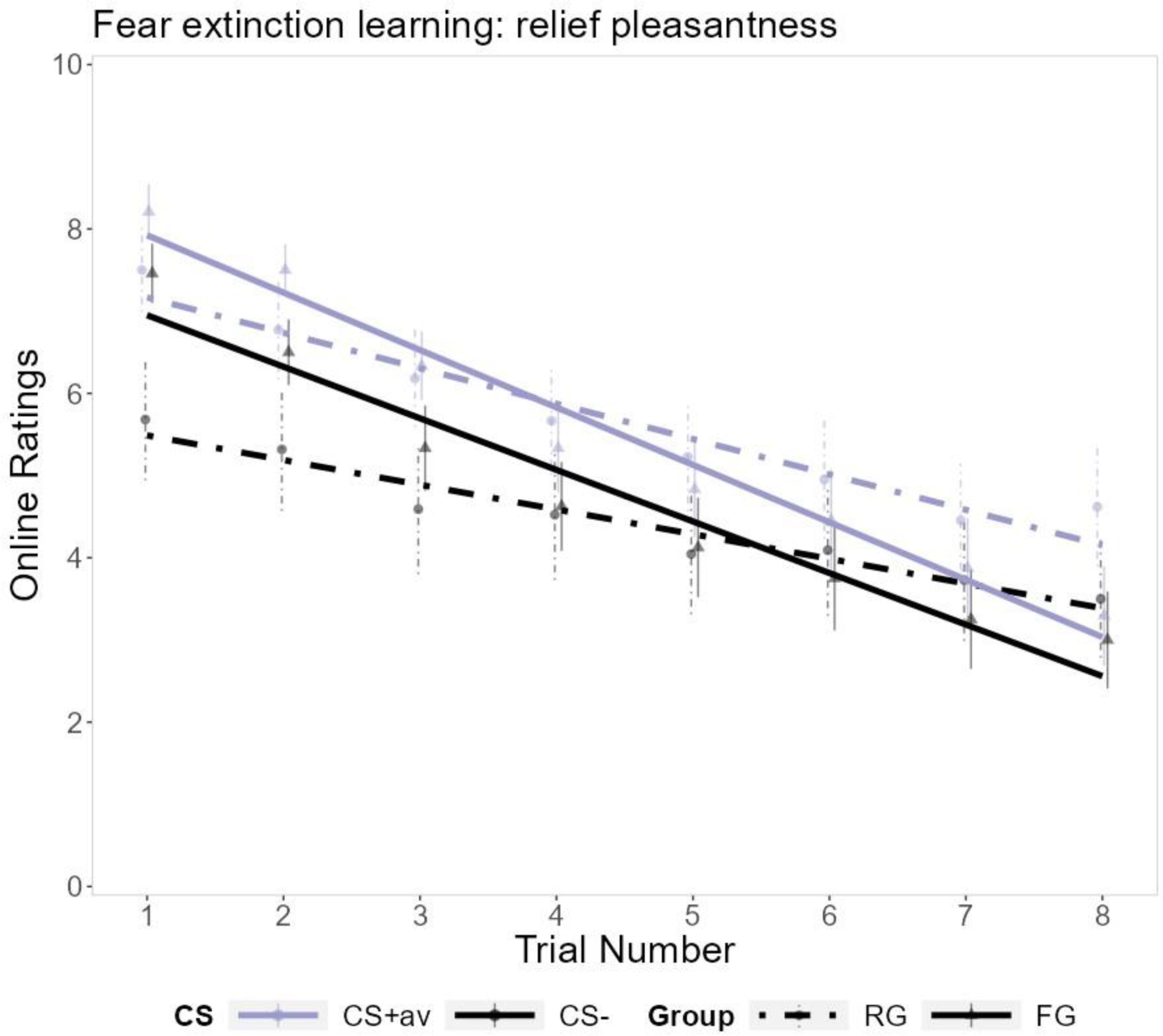
The relief pleasantness ratings for the Fasting Group (FG, continuous lines) and the Re-feeding Group (RG, dashed lines). Resembling the trajectory of a rPE, general relief significantly decreased over the trials (Trial: *β* = - 0.431 ± 0.083, df = 43.988, t = - 5.200, *p* < 0.001, 95% CIs [- 0.582:- 0.279], RESI = 0.757), and was significantly lower for the CS- (*β* = - 1.352 ± 0.144, df = 640.000, t = - 9.357, *p* < 0.001, 95% CIs [- 1.634:- 1.069], RESI = 0.442). Additionally, the Fasting Group showed a significantly faster decrease in relief than the Re-feeding Group. Error bars represent the standard error of the mean, while the lines represent the model fit for each CS.

#### Neural data

##### A priori hypothesis

###### Group-related difference during US omission (ROIs)

*Nac.* The results from the two sample t-test for the CS- versus CS+av contrast revealed a significantly smaller differential activation in the Fasting Group compared to the Re-feeding Group (t = - 2.672, df = 37.477, *p* = 0.005, 95% CIs [- inf:- 0.056], *d* = 0.803), **figure 5A**, right graph. The results from post hoc t-tests run separately for each CS showed that this difference was driven by the expected smaller response to the CS- in the Fasting Group compared to the Re-feeding Group (CS-: t = - 2.618, df = 39.347, *p* = 0.006, 95% CIs [- inf:- 0.037, *d* = 0.781]).

**Figure 5.**
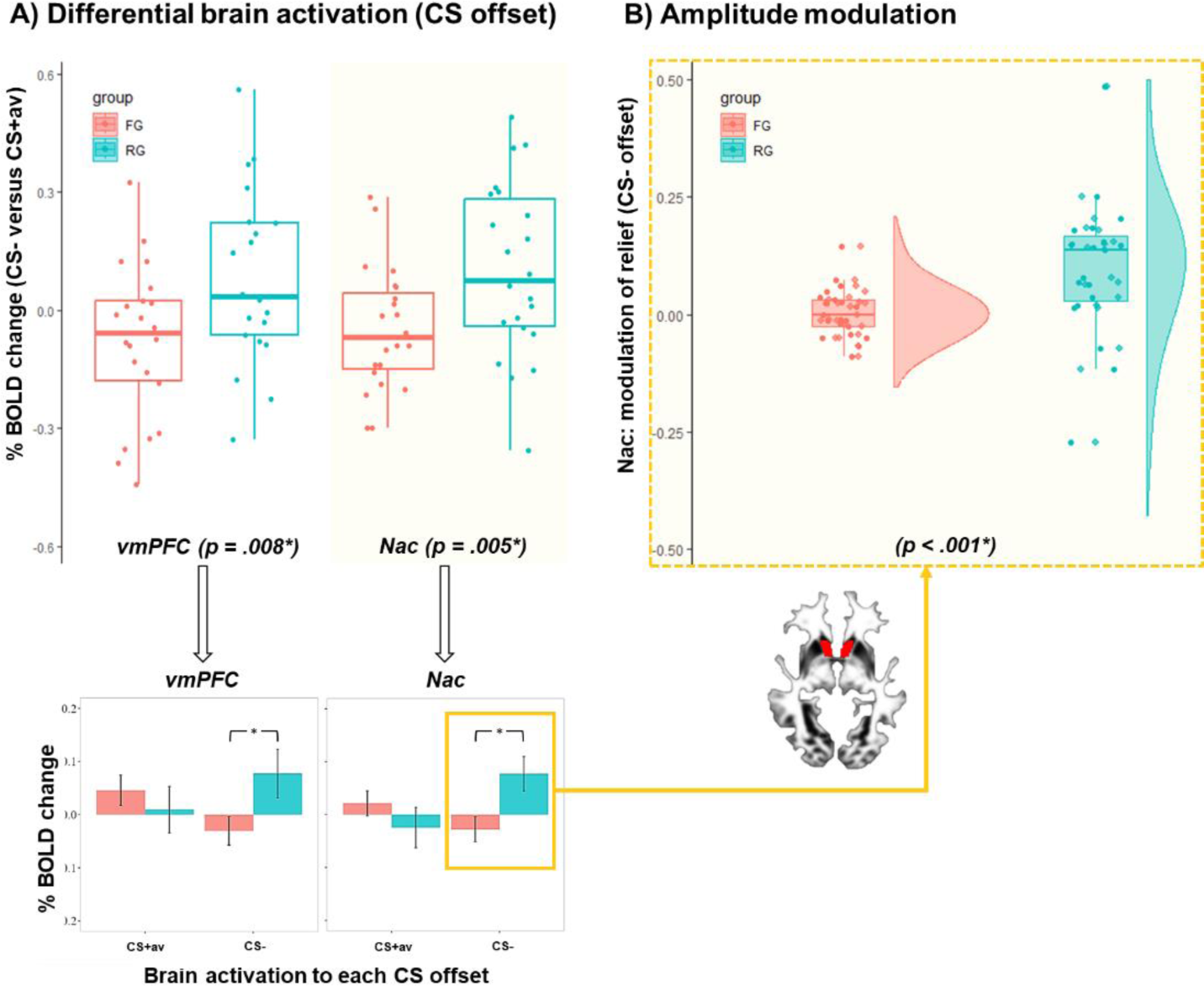
**A)** The percentage change in BOLD signal in the vmPFC *(left graph)* and Nac *(right graph)* in response to US omissions at the CS- versus CS_+av_ offsets for both the Fasting Group (FG, in orange) and the Re-feeding Group (RG, in blue). Compared to the Re-feeing Group, the Fasting Group showed a significantly smaller differential brain activation driven by the expected reduction in vmPFC and Nac activity in response to the CS-. **B)** The results from the amplitude modulation in AFNI showed a positive modulation of the relief rating on the activation of the Nac during the CS-offsets in the Re-feeding Group only. In the Fasting Group, the main activation for the CS-offset and the relief modulation approached a zero value.

*vmPFC.* The two samples t-test on the contrast (CS-versus CS+av) revealed the hypothesized smaller differential activation in the Fasting Group (t = - 2.506, df = 39.969, *p* = 0.008, 95% CIs [- inf:- 0.053], *d* = 0.767), see **figure 5A**, left graph. The results from post hoc t-tests run separately for each CS showed that this difference was driven by the expected smaller response to the CS- in the Fasting Group compared to the Re-feeding Group (CS-: t = - 2.027, df = 34.664, *p* = 0.025, 95% CIs [- inf:- 0.018], *d* = 0.611).

*VTA.* No significant group differences were found.

###### Amplitude modulation: the role of relief in US omission

In our fourth hypothesis, we examine if the group difference in brain activations found during the CS- offsets in the vmPFC and Nac were related to subjective relief on a trial-by-trial level using amplitude modulation analysis **(hypothesis 2-b)**.

*Nac*. Indeed, relief was positively related to Nac activations during the CS- offset across the two groups (t = 2.937, df = 36, *p* = 0.006, 95% CIs [0.013:0.072]). Furthermore, this relation differed significantly between the two groups (t = - 3.819, df = 21.331, *p* < 0.001, 95% CIs [- inf:- 0.055], *d* = 1.332), **figure 5B**. This group difference was driven by a significant positive modulation of relief only in the Re-feeding Group (t = 4.018, df = 16, *p* < 0.001, 95% CIs [0.046:0.147]), and not in the Fasting Group (*p* = 0.792).

*vmPFC.* Unexpectedly, no significant modulation was found, with or without accounting for group effects.

#### Skin Conductance Responses

In a secondary preregistered analysis, we investigated if fasting changed SCRs to US omissions. The results from the GLMM (logit component) showed a main effect of Group (omnibus: *ꭓ^2^* = 7.736, df= 1, *p* = 0.005). The results from post hoc analysis run separately for each CS indicated that compared to the Re-feeding Group, the Fasting Group showed lower probability in SC responding to US omission following both the CS- (*β* = - 1.090, ± .453, z = - 2.406, *p* = 0.016, 95% CIs [- 2.050:- 0.198], Exp-*β* = 1.188) and the CS+av (*β* = - 1.149, ± 0.444, z = - 2.585, *p* = 0.010, 95% CIs [- 2.095:- 0.289], Exp-*β* = 1.320). In an additional sensitivity analysis, we confirmed the same significant effect of fasting when the three participants who did not show any SCR during omission for any of the CSs were excluded. Results from the following LMM (lognormal component) showed no group-related effects.

We then added relief ratings to the GLMM model and found that in addition to the fasting intervention, relief was a significant positive predictor of the probability of SCRs (*β* = 1.147, ± 0.051, z = 2.851, *p* = 0.004, 95% CIs [0.047:0.256], Exp-*β* = 1.320). When we allowed relief ratings to interact with the factor Group, the group effect disappeared, while the interaction between Group and Relief was significant, with the Re-feeding Group showing a higher modulation of relief in the probability of SCRs (*β* = 0.176, ± 0.076, z = 2.315, *p* = 0.020, 95% CIs

[0.037:0.308], Exp-*β* = 0.030). This result suggests that fasting reduced the general probability of SC responding at CS offset, while higher relief ratings were associated with a higher probability of SC responding only in the Re-feeding Group, see **figure 6**. This result was similar to what we found at the brain level in the modulation analysis, although no specific for the CS-.

**Figure 6.**
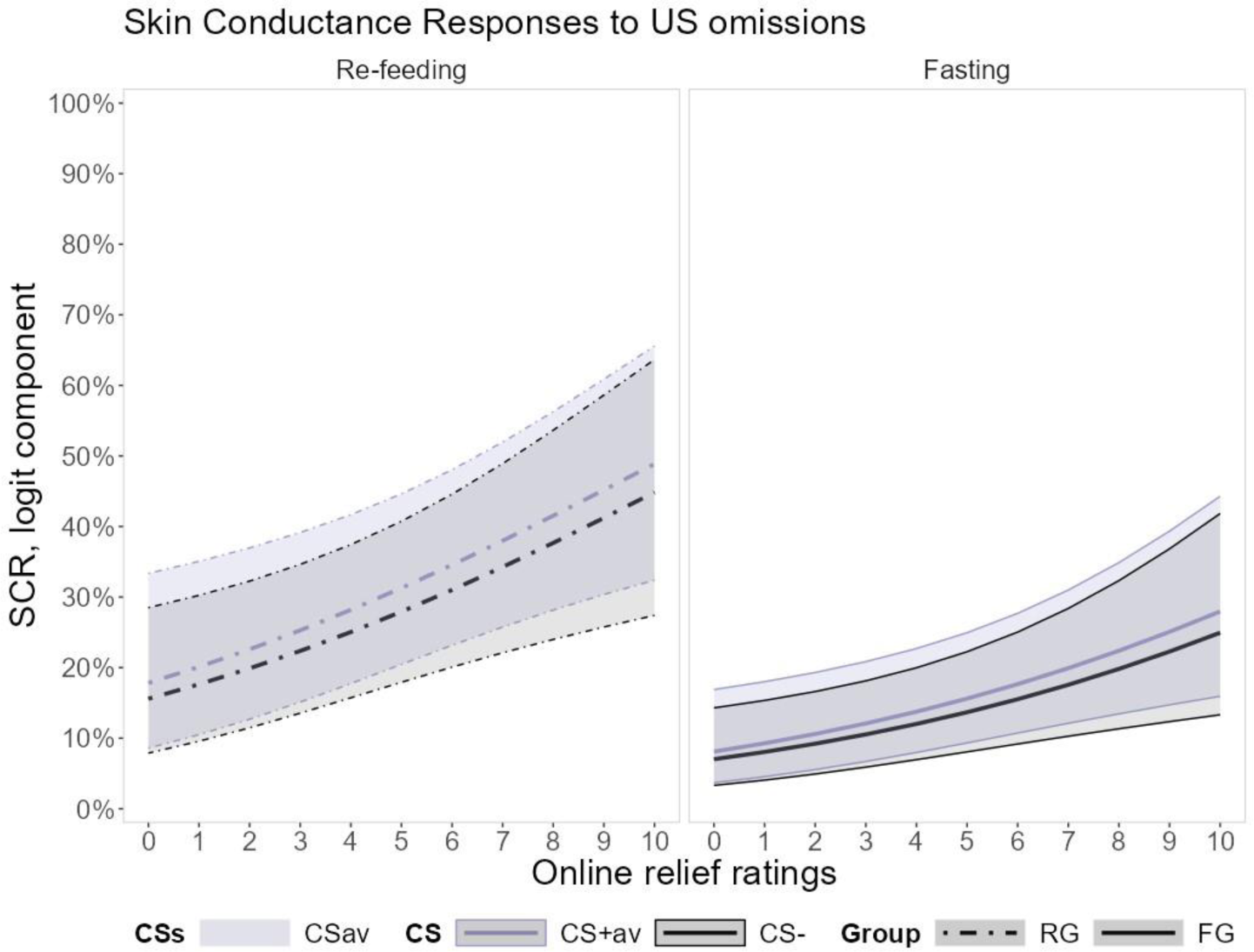
The probability of Skin Conductance responding (accumulated average over the trials) in the physiological signal that followed US omissions at the CS- (black) and CS+_av_ (blue) offsets, for both the Re-feeding Group *(left panel)* and the Fasting Group *(right panel).* The Fasting Group showed a minimization in the probability of responding, while in the Re-feeding Group the higher probability of responding was explained by higher relief ratings.

##### Explorative analysis: main effect of US omissions in other brain regions

For the avoidance and fear extinction learning phase, in the whole sample, we explored how other important PE-related areas of the brain responded to differential US omissions (CS+av versus CS-). For this purpose, instead of a whole-brain analysis as it was preregistered, we ran a voxel-based analysis using an extensive mask of interest (to reduce the problem of correction for multiple comparisons), see supplementary material for the areas included. In the supplementary material, we also report the ROIs analysis run in the whole sample.

###### Avoidance: voxel-based analysis within the extended mask

The results from a paired t-test between the two CSs (CS+av versus CS-) showed no significant differences. After a successful button press, the CS+av offset, similarly to the CS- offsets, elicited brain activity in the Putamen, Caudate (bilaterally), which extended to the Nac (left), and in area 32 of the vmPFC (which was an extension of a bigger cluster in the superior part of the Anterior Cingulate Cortex (ACC) and the dorsomedial PFC, dmPFC), see **table S3** in the supplementary material for further results, and **figure 7A**, for the general activation to US omissions (CS+av plus CS- versus baseline). No significant activation was found in VTA unless general US omission was compared to activation during baseline, see supplementary material.

**Figure 7.**
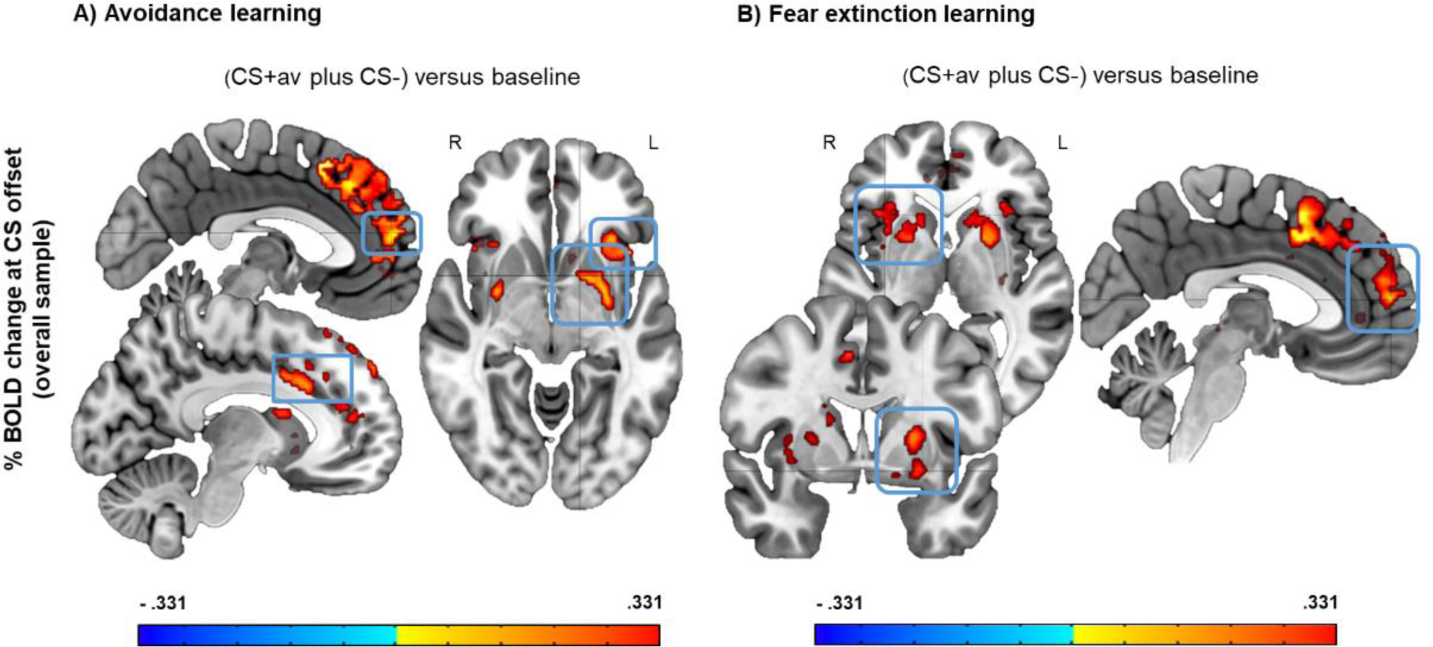
**A)** The general omission of the US (contrast: CS+_av_ plus CS- offsets versus baseline during avoidance learning), elicited significant brain activations similar to those reported in **Table S3** for each separated CS vs baseline. Unexpectedly, we did not find activation of the most orbital part of the vmPFC but rather a significant activation in the a32sg of the left vmPFC (peak: x: 2.5, y: -53.5, z: 0.7) which included also the dACC (*sagittal sections*). There was also a significant activation of the Caudate and Putamen (bilaterally) which activations on the left (peak: x: -28.5, y: -2.5, z: -8.1) extended to the left Nac (*axial section)*, (significant for a cluster extended (FWE-corrected) threshold alpha below 0.05 with an initial *p* = 0.001 uncorrected). All the brain activations mentioned were significant independent of how many times the participants pressed the button for the CS-. **B)** Shows the brain activity in response to general US omissions during the fear extinction learning phase: CS+_av_ plus CS- offsets versus baseline. The general omissions of the US elicited significant brain activations similar to those reported in **Table S4** for each separated CS vs baseline. We found significant activations in the Caudate and Putamen (bilaterally) (left peak: x: 28.5, y: 2.5, z: -8.1) (*axial and coronal sections),* in the superior part of the a32 vmPFC (peak: x: -1.5, y: -57.5, z: 11.7) (*sagittal section)* which included also the dACC, and significant activation of the Insule (right peak: x: -35.5, y: -17.5, z: 5.1). The results were significant for a cluster extended (FWE-corrected) threshold alpha below 0.05 with an initial *p* = 0.001 uncorrected.

###### Fear extinction: voxel-based analysis within the extended mask

The results from a paired t-test showed that the CS+av versus CS-contrast did not yield any significant result. This is not surprising since in a new extinction context, high relief levels have been found for both the CS- and the CS+av^6,8^ offsets, while the vmPFC and Nac respond similarly to omissions after CS- and CS+ (see discussion in^27,29^ and^20^ for Nac). We found that the CS+av as well as the CS- (vs the baseline) elicited similar brain activity during US anticipations (see supplementary material), and also to US omissions in the Putamen and Caudate (bilaterally), which cluster in the Left Putamen marginally extended to the Nac, and in the superior area 32 of the vmPFC, which was an extension of a bigger cluster in the superior part of dorsal Anterior Cingulate Cortex (dACC) and the dorsomedial PFC (dmPFC), similarly to the clusters found in the avoidance phase. These results have to be interpreted cautiously since these activations might include the response to the appearance of the black screen (CS offset). See **table S4** in the supplementary material for further results, and **figure 7B**, for the general activations to US omissions (CS+av plus CS-versus baseline).

## Discussion

The aim of this study was to investigate the neural mechanisms by which overnight fasting decreases unnecessary/ineffective avoidance and speeds up fear extinction learning with a specific focus on the reward system (Nac, vmPFC, and VTA) and the relation with subjective relief experiences. For this purpose, we replicated our previous experiment ^5^ in the fMRI scanner, comparing two groups that fasted 14-16 hours overnight and consumed (standard) breakfast before (Re-feeding Group) or right after the experiment (Fasting Group).

As in our previous study, overnight fasting led to more discrimination between the avoidance responses to the CS+av (when the US was actually avoidable, which is an adaptive behavior), and the CS+unav (when avoidance was ineffective) or the CS- (when avoidance was unnecessary). This time, however, this optimization was driven by increased effective avoidance to the CS+av, rather than decreased avoidance to the CS+unav and CS- as in our previous experiment. Against our expectations, we did not find fasting effects on neural or skin conductance responses to US omission that were of statistical relevance (although they pointed to the hypothesized direction), while the absence of any effect of fasting on US omission-induced relief did not allow a further investigation of relief as a potential modulation on the avoidance behavior, as we previously showed.

During subsequent fear extinction, in which CS+av and CS- were presented in a new context without the avoidance button and without the US, subjective relief during US omissions decreased at a faster rate and minimized more in their residual levels in the Fasting Group than in the Re-feeding Group (for both CS+av and CS-), in line with our previous findings^5^. This finding broadly supports the work of two studies in rodents which investigated the effect of acute fasting on fear extinction learning. Verma et al. found that acute fasting decreases the consolidation of fear when applied before acquisition, facilitates fear extinction when applied after conditioning and before extinction, and increases extinction recall after re-feeding when applied before extinction learning^4^. Similarly, Huang et al. found that acute fasting enhanced fear extinction and its retention^3^.

In male humans, only one behavioral study ^30^ investigated the effects of overnight fasting on fear extinction and its retention. The authors found that the retention of extinguished fears (but not learning) is enhanced by an overnight fasting procedure applied before fear acquisition. In the same vein as Shi et al., we found no effect of overnight fasting in anticipatory SCRs during fear extinction. However, we found an effect on PE-related outcome measures which were not investigated by Shi et al. The differences between our and Shi et al. results might also depend on the protocols used (one day protocol for our fMRI study, and a two days protocol for Shi et al.), and on gender.

We showed here for the first time that US omissions elicited lower vmPFC and Nac activations in the Fasting Group compared to the Re-feeding Group (when contrasting CS-offsets to CS+av offsets). Moreover, Nac activations and general SCRs were positively modulated by subjective relief in the Re-feeding Group, but not in the Fasting Group, where those activations approached a minimal value. This suggests again that overnight fasting speeds up fear extinction through a reduction in responsivity to/pleasantness of US omissions.

The Nac exerts a ‘hub’ function integrating online PEs with reward pleasantness in classical reward-based learning^31^. Here, the Nac might exert a similar function, by integrating PEs from threat omissions with their hedonic impact (relief ‘liking’). Overnight fasting minimizes the modulation of this hedonic component as well as online PEs in Nac responses to the safest circumstance (CS- offset) and not to dangerous cues (CS+). In a fed state, instead, the magnitude of these rPEs in the Nac increases with the amount of pleasantness of the relief experienced in response to the omissions of the safe CS. This evidence further supports our theory that, while fasting increases rPEs in appetitive learning, it exerts a reverse effect on rPEs that govern fear extinction learning (at least for what concerns the CS-). Now, in a laboratory setting, changes in the responses to threat omissions signaled by the safe cue are often of less interest. In the context of clinical anxiety, however, we often observe increased avoidance and relief for the safe CS^32^. Hence, how results suggest that overnight fasting might be a useful adjunct to psychotherapy (e.g. exposure).

Similarly to what was found in the Nac, overnight fasting minimizes the response of the vmPFC to the CS-. The vmPFC, is known to be central in decoding the value of a CS^33^, and, in concert with the Nac, shows a decrease in response to food when food is devaluated via satiety^34–36^. Hence, we interpret the present fasting-induced reduction in vmPFC at the offset of the CS- as a potential fasting-induced reduction in the value of safety, in line with our theory^5^. As for the Nac, we expected to find a modulation on a trial-by-trial level with relief also in the vmPFC. This was however not the case and in line with the results from a previous fMRI study investigating the neural correlates of relief within a pain omission paradigm. Specifically, Leknes and colleagues^37^ found a similar activation in vmPFC and Nac when comparing brain activations in response to classical rewards and unexpected omissions of painful stimulations. In their study, the vmPFC and the Nac were found active during (pain) omissions, but only the response of the Nac was positively related to relief ratings. Our paradigm deviates from that of Leknes especially in terms of its learning nature. It seems therefore that differently from the Nac, the vmPFC during learning does not compute the pleasantness of the online deviations from previous US omissions. Hence, our results show that overnight fasting has opposite effects on rPEs within the appetitive and fear extinction context. It remains to be elucidated if these opposite effects on rPE signaling is dopamine-dependent.

Previous animal research has established the role of prolonged fasting and food restriction in increasing midbrain dopaminergic response to rewarding stimuli^38–41^. Specifically, fasting increases the salience and reward value of food in VTA-Nac reward circuitry^14^, and triggers motivated appetitive behaviors upon the activation of both the (mesolimbic) ‘wanting’ and the (interceptive) ‘needing’ systems^42^. When acute, fasting also increases dopaminergic turnover in response to salient events. This increase might depend on a reduction of midbrain dopaminergic activity during rest^41^ or on changes in the striatal opamine reuptake transporter (DAT)^39^. In either case, acute fasting ultimately facilitates the response of midbrain dopaminergic neurons that are central to reward and safety learning^15,16^. Hence, our results might depend on the effects of lower tonic DA on leaving more room for a phasic dopaminergic burst if a reward is consumed, and/or on a reduced tonic DA itself. To address this question, studies with similar protocols could be used together with pharmacological manipulations of the dopaminergic system or Positron Emission Tomography using appropriate radioligands.

To sum up, in healthy female individuals, overnight fasting improves fear extinction learning by reducing relief pleasantness, reward-related brain areas, and electrodermal responses during threat omissions/PEs.

## Limitations

The absence of the expected effect of fasting in decreasing unnecessary and ineffective avoidance might be explained by the change in the experimental environment. The MRI environment, compared to a regular lab setting, undoubtedly represents a more stressful context ^43^. Stress, on top of an overnight fasting procedure, might affect avoidance learning differently from only a fasting procedure. This might be because avoidance learning involves the activation of the motivational system and behavioral engagement. The passive process of fear extinction learning might be more robust to this type of additive effect of stress and fasting instead. To test this hypothesis, it would be important to investigate the effects of stress procedures (and biomarkers of stress) in our fasting experiment.

Second, the effects of fasting were present across all three measurements of US omission, although related to time only in the relief ratings and stimulus-specific only at the brain level, while neither to time nor CSs in the SCR. These specificities in each type of measurement remain difficult to explain at the moment. Speculatively, the absence of trial/time-related effects at the brain level can be explained by the single-trial-based learning nature of the ART, which aspects were better captured by the individual relief ratings instead. To overcome this limit, future versions of the ART could include a partial reinforcement schedule. Nonetheless, as already recommended in a previous commentary^44^, we attempt this analysis to also bring more awareness to the importance of trial/time effects in learning paradigms used in neuroimaging and behavioral studies, which are currently largely neglected. Crucially, averages-based comparisons should be limited since they might hide crucial information in how learning generates and evolves.

Differently, the presence of a stimulus-specific effect of overnight fasting only at the brain level might specifically depend on the better ability of the MRI (compared to cognitive ratings and peripheral changes in sudoriparous glands) to capture online rPE.

Next, replications of this study should take into account the difficulty to find a significant discrimination in brain responses (as well as in other measures) to US omissions signaled by the two CSs when fear extinction takes place in a new context. In this case, indeed, we often observe an increase in US expectation/relief for the safe CS-. That is why differences between the two types of US omissions might not be (significantly) detectable, unless, as in our case, the homeostatic state is manipulated. Replication of our study should also investigate whether these effects of fasting hold if fasting is applied after fear conditioning and when fear extinction is performed in the conditioning context. Furthermore, we did not find group differences in VTA. The small size of such brain region (volume of 360ml) makes it difficult to capture its change in activity by a whole-brain slicing sequence during one trial-based learning, see^21^.

## Conclusion

The current results highlight the importance of an investigation of overnight fasting in a clinical population, as well as the necessity to take into account the homeostatic state of the participants in neuroimaging studies that investigate safety learning.

## Methods

### Participants

Fifty healthy Dutch females (age: 22.21/3.53, Body Mass Index [BMI]: 21.26/2.08, Mean/SD) participated in the study after providing written informed consent. The study was approved by the local Medical Ethics Committee (EC). Exclusion and inclusion criteria, compensation, as well as the procedures for the calculation of the sample size, are reported in the supplementary material.

### Procedures

#### Overnight fasting manipulation

The fasting procedure was identical to our previous behavioral study^5^. Briefly, participants underwent a14-16h period of fasting, starting from the evening before the fMRI session (which took place in the morning). During the fasting period, only consumption of water was allowed. Before the fMRI experiment, the Re-feeding Group (RG) consumed a standard breakfast upon arrival in the lab, while the Fasting Group (FG) received the same breakfast only after the fMRI experiment. Differently from the previous experiment, the start of the fasting period (set at 6 PM) was individually adjusted depending on scanning availability the next day (start of the scanning session was always between 9:00 and 11:30 AM).

#### Visits and fMRI examination

The testing procedures were similar as in our previous study with a few exceptions. These involved the inclusion of a neuropsychological assessment to exclude the effects of fasting on working memory, selective attention and cognitive efforts, and removal of one of the test phases of the ART due to time limitation, see **figure 1** for details.

### Apparatus

#### Stimuli

Visual stimuli as well as the procedures and the delivery of the electrical stimulation were the same as those reported in our behavioral study.

Briefly, two different contexts were presented on the computer screen (an office room and a conference room). Both contexts contained a desk lamp that could light-up in three different colors: red, blue, and yellow, which colors served as conditional stimuli (CSs). We used an electrical stimulation as the unconditional stimulus (US). The CSs+ were followed by the US while the CS- was not. Avoidance action cancelled the electrical stimulation after the CS+avoidable, but not after the CS+unavoidable, CS+unav. In all trials, we used a jittered presentation of the stimuli to avoid learning effects, see **figure 1**.

The electric stimulation consisted of high-voltage brief 2-ms electrical pulses delivered at the forearm of the non-dominant hand of the participant. The level of the electrical stimulation was selected by the participant before the start of the task via a gradual intensifying procedure.

Ketone levels were measured using Ketostiks strips (Ascensia Diabetes Care), while glucose levels were assessed using a standard glucometer device (GlucoMen® areo Sensor). Skin conductance levels (SCL) were measured using an fMRI-compatible Biopak set-up, which was linked to two sensors attached to the palm of the left hand of the participant.

#### Self-report ratings

*Physical* (‘How hungry are you?’) and *emotional* (‘How strong is your desire to eat?’) hunger was measured on a Likert scale ranging from 0 (’Not at all’) to 10 (’Very hungry’) before and after the diet manipulation. *Retrospective US expectancy* for each stimulus was measured on a Likert scale ranging from 0 (’Shock not expected/not afraid at all’) to 10 (’Shock expected/very much afraid’) at the end of each learning phase of the task. *Relief pleasantness* was measured every time the US was not delivered on a Likert scale ranging from 0 (’Neutral’) to 10 (’Very pleasant’) appearing on the computer screen. *Motivation to avoid* (’How much would you like to avoid the electrical stimulation’) was measured on a similar scale ranging from 0 (’Not at all’) to 10 (’Very much’) appearing on the computer screen at the end of each learning phase.

#### Behavioral outcome

Avoidance actions were measured in terms of button presses.

### Paradigm

A full description of the ART task can be found in^5^. Briefly, during the *Pavlovian phase*, three light colors presented in context A (office room) are used as CSs that signal the absence or the presence of an upcoming unpleasant electrical stimulation (used as US) to the non-dominant forearm: the CS+av and the CS+unav are both followed by the US, while the CS- is not. Next, during the *avoidance learning phase*, which takes place against the same room picture (context A), a red button appears during all CS. Clicking the button effectively prevents the US during CS+av, but not during CS+unav. Finally, in the *fear extinction phase* with response prevention, the context changes from an office to a study room (context A◊B) and the CS+av and CS- are presented without the presence of the red buttons and without any US. A relief rating scale is presented every time the US is omitted. Retrospective US-expectancy ratings are collected at the end of each learning phase.

### Analysis plan

#### General analysis

For each measure, specific exclusion criteria are reported in the supplementary material. All the variables of interest were analyzed using (generalized) linear mixed models (GLMM and LMM)^45^ in R-studio (Version 1.4.1717©2009-2021, RStudio) and AFNI (22.0.04) (3dDeconvolve/3dREMLfit). Before reporting the planned contrasts of interest, we also reported any Group-related effect from an omnibus test (using car::anova(glmer) in rstudio with a type III Wald test for ꭓ^2^ GLMM, or by using anova(lmer) for LMM). The independent variables were ‘Group’, type of ‘CS’, and ‘Trial’. ‘Trial’ was used as a categorical dichotomous variable for the analysis of the retrospective US expectancy ratings [first and last presentation of each CS], and as a continuous de-meaned variable for the remaining analysis. For each measure, the planned contrast of interest refer to the Group by CS interaction. We ran post hoc Bonferroni corrected tests for any other eventual Group or Group by Trial effect (for which we did not have a priory hypothesis). We reported the 95% confidence interval (CI) for the estimate of each GLMM/LMM result of interest [lower: upper]. For the GLMMs, as a measure of effect size, we use the exponentiated estimates (Exp-*β*) (with Exp-*β* >1 indicating increased occurrence of an event); for the LMMs, we used the Robust Effect Size Index, RESI, rstudio: resi_pe^46^ (with [0, 0.1] = small effect, [.1, 0.25] = small to medium effect, [0.25, 0.4] = medium to large effect, and large effects greater than 0.4).

For the brain data, we were primarily interested in Groups differences for the CS- vs CS+av contrast. For this investigations we used two sample t-tests and the Cohen’s *d* to quantify effects sizes (0.2 = small effect, 0.5 = moderate effect, 0.8 = large effect). We also included explorative analysis on the effect of time (early/late learning) and trial (time series analysis which can be intended as a growth curve modeling). For the first type of time-related analysis, we first run a t-test on early compared to late omissions (CS- versus CS+av)early versus (CS- versus CS+av)late (average of the beta weights calculated on the first and last four trials for each CS). For the second type of analysis, we run a LMM for each ROI by including ‘Trial’ as covariate of interest. In general, since the previous effects of fasting were found mostly on the CS-, the CS+av was here used as the reference for comparisons in the t-tests, LMMs and GLMMs. Within each LMM model, the normality of the distribution of the residuals was visually checked and corrected with a log-transformation if needed. In all the GLMM and LMM models we included random subject, random slope by subject, random CS by subject, and random slope by CS by subject only when these terms did improve the model’s fit based on the Akaike’s information criterion (AIC). Significance was set at a *p* < 0.050.

### Skin Conductance Responses

### SCR preprocessing

A bandpass filter^47^ was applied to the SCL data. These were then analyzed in MATLAB R2018b to calculate the skin conductance responses (SCRs) for the onset and offset of the CS. Each SCR was calculated by subtracting the average SCL during the 1 second prior to appearance of the CS on the screen (US-anticipation), to the peak found during the presentation of the CS (7.5-9 sec). A z-transformation was applied to each value to increase the comparability of the results. We also changed the threshold applied to the responses from 0.02 to 0.01 to reduce the number of non-responders.

### SCR analysis

After the removal of the outliers (IQR, 1.5 within-subject on a trial level) we run a mixed-effects mixed distribution model for zero-inflated data^48^. For SCRs which ln transformation did not resulted in a normal distribution, we run a mixed-effects mixed distribution model for zero-inflated data (for the logistic component, we ran a GLMM while for the lognormal component we used a LMM analysis), see our previous paper for further details.

### MRI data

### MRI preprocessing

#### - Anatomical MRI data preprocessing

A total of 50 T1-weighted (T1w, 1mmx1mmx1mm) images were found within the input BIDS dataset. The T1-weighted (T1w) image was corrected for intensity non-uniformity (INU) with N4BiasFieldCorrection ^49^, distributed with ANTs 2.3.3 (RRID:SCR_004757)^50^, and used as T1w-reference throughout the workflow. The T1w-reference was then skullstripped with a *Nipype* implementation of the antsBrainExtraction.sh workflow (from ANTs), using OASIS30ANTs as target template. Brain tissue segmentation of cerebrospinal fluid (CSF), white-matter (WM) and gray-matter (GM) was performed on the brain-extracted T1w using fast (FSL 5.0.9, RRID:SCR_002823)^51^. Volume- based spatial normalization to two standard spaces (MNI152NLin2009cAsym, MNI152NLin6Asym) was performed through nonlinear registration with antsRegistration (ANTs 2.3.3), using brain- extracted versions of both T1w reference and the T1w template. The following templates were selected for spatial normalization: *ICBM 152 Nonlinear Asymmetrical template version 2009c* [^52^, RRID:SCR_008796; TemplateFlow ID], *FSL’s MNI ICBM 152 non-linear 6th Generation Asymmetric Average Brain Stereotaxic Registration Model* [^53^RRID:SCR_002823; TemplateFlow ID: MNI152NLin6Asym].

#### - Functional MRI data preprocessing

For each of the 3 BOLD runs found per subject (across all tasks and sessions), the following preprocessing was performed. First, a reference volume and its skull-stripped version were generated using a custom methodology of *fMRIPrep*. Susceptibility distortion correction (SDC) was omitted. The BOLD reference was then coregistered to the T1w reference using flirt (FSL 5.0.9, Jenkinson and Smith, 2001) with the boundary-based registration (Greve and Fischl, 2009) cost-function. Co- registration was configured with nine degrees of freedom to account for distortions remaining in the BOLD reference. Head-motion parameters with respect to the BOLD reference (transformation matrices, and six corresponding rotation and translation parameters) are estimated before any spatio-temporal filtering using mcflirt (FSL 5.0.9, Jenkinson et al., 2002). BOLD runs were slice-time corrected using 3dTshift from AFNI 20160207 (^54^, RRID:SCR_005927). The BOLD time-series (including slice-timing correction when applied) were resampled onto their original, native space by applying the transforms to correct for head-motion. These resampled BOLD time-series will be referred to as *preprocessed BOLD in original space*, or just *preprocessed BOLD*. The BOLD time- series (with 2mmx2mmx2mm voxel size) were resampled into standard space, generating a *preprocessed BOLD run in MNI152NLin2009cAsym space*. First, a reference volume and its skull- stripped version were generated using a custom methodology of *fMRIPrep*. Several confounding time-series were calculated based on the *preprocessed BOLD*: framewise displacement (FD), DVARS and three region-wise global signals.

FD was computed using two formulations following Power (absolute sum of relative motions^55^,,and relative rootmean-square displacement between affines^56^. FD and DVARS are calculated for each functional run, both using their implementations in *Nipype* (following the definitions by^55^). The three global signals are extracted within the CSF, the WM, and the whole-brain masks. Additionally, a set of physiological regressors were extracted to allow for component-based noise correction (*CompCor*^57^). Principal components are estimated after high-pass filtering the *preprocessed BOLD* time-series (using a discrete cosine filter with 128s cut-off) for the two *CompCor* variants: temporal (tCompCor) and anatomical (aCompCor). tCompCor components are then calculated from the top 2% variable voxels within the brain mask. For aCompCor, three probabilistic masks (CSF, WM and combined CSF+WM) are generated in anatomical space. The implementation differs from that of Behzadi et al. in that instead of eroding the masks by 2 pixels on BOLD space, the aCompCor masks are subtracted a mask of pixels that likely contain a volume fraction of GM. This mask is obtained by thresholding the corresponding partial volume map at 0.05, and it ensures components are not extracted from voxels containing a minimal fraction of GM. Finally, these masks are resampled into BOLD space and binarized by thresholding at .99 (as in the original implementation). Components are also calculated separately within the WM and CSF masks. For each CompCor decomposition, the *k* components with the largest singular values are retained, such that the retained components’ time series are sufficient to explain 50 percent of variance across the nuisance mask (CSF, WM, combined, or temporal). The remaining components are dropped from consideration. The head- motion estimates calculated in the correction step were also placed within the corresponding confounds file. The confound time series derived from head motion estimates and global signals were expanded with the inclusion of temporal derivatives and quadratic terms for each^58^. Frames that exceeded a threshold of 0.9 mm FD or 2.0 standardised DVARS were annotated as motion outliers (spikes).

Participants with more than 15% of spikes were excluded from the extinction learning phases, while a threshold of 20% of spikes was applied to the avoidance phase (since this last phase lasts 15 minutes and includes button presses). All resamplings can be performed with *a single interpolation step* by composing all the pertinent transformations (i.e. head-motion transform matrices, susceptibility distortion correction when available, and co-registrations to anatomical and output spaces). Gridded (volumetric) resamplings were performed using antsApplyTransforms (ANTs), configured with Lanczos interpolation to minimize the smoothing effects of other kernels^59^. Non-gridded (surface) resamplings were performed using mri_vol2surf (FreeSurfer). We included 32 regressors to account for noise-movement related effects: 24 motion parameters that include the 6 rigid-body parameters for rotation (3) and translation (3) and 2 parameters for WM and CFS, plus their first-order associated derivatives and their square terms. We did not include global signal among these regressors. A smoothing of 4 4 4 mm was applied to the data. The images were then scaled, so that each voxel had a mean signal intensity of 100. This produces small signal detectable on a trial by trial level, but allows us to correctly compare the two groups by specifying any change relative to the mean as percent signal change.

#### - Selection of the Regions Of Interest (ROIs)

The vmPFC mask was obtained from AFNI (area 14/32/24, bilateral). Since we consider the pleasantness of relief from omission of a threat a type of reward, the selection was made based on an activation likelihood estimation (ALE) meta-analysis comparing the brain responses to monetary, erotic and food reward outcomes^60^.

The Nac and VTA masks were obtained from the Canlab data-repository (http://canlab.github.io), while the masks for the sub-nuclei of the Nac (Shell and Core) were obtained from the brainnetome atlas (https://atlas.brainnetome.org). For the analyses on the main effect of the task (e.g. brain response to US anticipation/omissions across groups), we used a ROIs analysis (averaged beta weights), as well as a voxel-based analysis on an extended mask that included our original vmPFC, Nac, VTA, and other rPE-related and reward-related brain areas, see supplementary material for details. Each mask was resampled to the original dimension of the functional images, in line with the MNI 2mmx2mmx2mm voxel size template.

#### - fMRI analysis

Since ART is a new paradigm in neuroimaging studies, after having addressed our primary research questions, we explore the main effects of the task during US omissions using t-tests at level of the ROIs, as well as a voxel-based analysis using the extended mask of interest.

The analyses were run with the 3dttest++ in combination with 3dClustSim in AFNI. For the results, we used a cluster-base correction for false positive with AFNI -3dClustSim (initial *p* = 0.001 and extended threshold alpha (FWE-corrected) of 0.05). A correction for a factor of 3 (VTA, Nac, and vmPFC) was applied to the alpha-value of the results from the t-tests/LMM for each ROI analysis (corrected with a factor of 3, *p* < 0.017).

#### - First-level analysis

The individual smoothed and scaled brain data from the avoidance and extinction learning phase of the ART were used to extract beta averages for the t-test analyses and to obtain the time-series beta weights for the LMMs.

The ROI analysis was done by using 3dDeconvolve to create a regressors matrix design and by using 3dREMLfit to run a GLM for the average BOLD signal across voxels of the ROI. A correction for noise correlation (ARMA) was applied at the trial-by-trial level. We repeated this process for the three ROIs. This allowed obtaining the beta estimates for the analysis of the early-late brain activations, as well as for the time series brain activations (on a trial-by-trial level). For both the avoidance phase (hypothesis 1) and the extinction learning phase (hypothesis 2) we were interested in the differential brain response at the time of the omission of the electrical stimulation during safety (CS-) and during CS+av offset (CS- versus CS+av). For the avoidance phase, the other regressors consisted of the activations during US anticipation (calculated from the disappearance of the red button till the end of the CS), brief activation during the button press, and activation during the time of the relief ratings. For the Pavlovian and extinction phases, the other regressors consisted of brain activation during CS presentations (US anticipation) and during the time of the relief rating.

#### - Second-level analysis

The results from the contrasts of interest from the first level analysis (CS- versus CS+av) were used within the second-level design to test for group-related differences (independent samples t-tests). When group-related effects were significant, an amplitude modulation analysis was done in AFNI by adding relief ratings as a mean-centered modulator for the brain activity of interest. In the explorative time-related analyses, besides the (CS- versus CS+av)early versus (CS- versus CS+av)late contrast, we implemented the time-series of the betas values in a Group by CS by Trial LMM analysis.

## Data availability

Original data are stored and secured within the KU Leuven facilities given ethical reasons. Processed anonymous data are available on the Open Science Framework (OSF), osf.io/yx8bq.

## Acknowledgements

BV was supported by a KU Leuven starting grant STG-18-00299, LVO is an associate research professor funded by the KU Leuven Special Research Fund (BOF), TB was supported by Consolidator Grant 648176 of the European Research Council. Preparation of this manuscript was also supported by KU Leuven research grant number C16/19/002 awarded to TB and BV and by a FWO project (G078920N) awarded by BV.

The funders had no role in study design, data collection and analysis, decision to publish or preparation of the manuscript.

The authors would like to acknowledge the support for the MRI acquisition by Dr. Ronald Peeters (MR physicist at the UZLeuven), and to all the participants.

## Contributions

BV, LVO and SP developed the concepts of the study and study design; SP performed the collection of the data, the data analysis and interpretation under the supervision of BV, TB and LVO; SP drafted the manuscript under the supervision of BV. All the authors provided revisions, conceptual and statistical feedback, and approved the final version of the manuscript for submission.

## Notes

https://archive.org/details/osf-registrations-yx8bq-v1

### Competing Interest Statement

The authors have declared no competing interest.

