## Supplementary for "Overnight fasting facilitates safety learning by changing the neurophysiological response to relief from threat omission"

**Supplementary Material**

**(Papalini et al.)**

### Exclusion and inclusion criteria

Exclusion criteria: current neurological (e.g. Epilepsia), respiratory, cardiovascular, metabolic, gastrointestinal, endocrine, renal or urinary diseases, psychiatric disorders, or other relevant medical histories; hypertension; lack of appetite; current or recent regular medication use (excluding contraceptives); smoking; high caffeine intake (> 1000 ml coffee daily or equivalent); Daily intake of soft drinks > 1000 ml; being pregnant or lactating; alcohol intake greater than 14 alcoholic units per week (one alcoholic unit = 10 gr ethanol); history of cannabis use or any other drug of abuse during the 3 months prior to the study; food or drug allergies; medical advice to avoid stressful situations; electronic implants (e.g., pacemaker); pain or other condition of the hand or the wrist; contraindications for the MRI exam; and currently following a fasting or restrictive diet. We also did not include individuals who used to skip breakfast.

We included only females (18-40-year old) taking hormonal contraceptives; right-handed and Dutch-speaking participants.

### Sample size calculation

The sample size was calculated based on the Group by CS interaction term of interest of our previous behavioral study[1]. For the calculation, we used General Linear Mixed Model Power and Sample Size (GLMMPSE) software (https://glimmpse.samplesizeshop.org/). For the computation, we use avoidance actions (button press) as the outcome variable. As a between-subjects factor, we included Group (Fasting and Fe-feeding group), and as a within-subjects factor we included CS (CS-, CS+avoidable, and CS+unavoidable). We selected a Target power of 0.8 and a Type I error rate of .05. The mean for the fasting group was .17 (CS-), .86 (CS+avoidable), and .36 (CS+unavoidable). The mean for the re-feeding group was 0.42 (CS-), .89 (CS+avoidable), and .5 (CS+unavoidable). The total standard deviation for the avoidance actions was .499, while the standard deviation for each CS was: 0.456 (CS-), 0.331 (CS+avoidable), and .495 (CS+unavoidable). The provided sample size (46 participants with a variability scale factor of .7) was increased to 50 to count for dropouts and potential (expected) bad recordings.

### Compensation

Each participant received monetary or a credits-based compensation based on the criteria accepted by the standard guideline of the Faculty of Psychology and the MEC.

### Exclusion from data analysis: number and reasons

One participant was excluded from the overall analysis of the avoidance phase since she did not perform the task (less than five clicks on the button). For the extinction phase, we included twenty-four participants in the FG and twenty-two participants in the RG (for a total of forty-six participants) in the ratings/brain-related data analysis. This exclusion was based on the presence of several spikes artifact in the MRI signal and/or on the impossibility to perform the fear extinction phase (time issues at the MRI facility).

In the analysis for the SCR (avoidance phase), forty-one participants were included since we excluded participants who showed bad recording in MRI and SCL acquisition. For the extinction phase, forty-four participants were included since three participants (in addition to those excluded from the brain data analysis) showed bad recordings in SCL.

**Construction of the extended mask**

The extended mask was used for validation purposes of the ART, to allow a more comprehensive investigation of several other potential rPE and emotional-learning-related brain areas, as well as to reduce the problem of multiple corrections. The mask included the original ROIs and the PeriAqueductal Gray, Amygdala, Hippocampus, Thalamus, dorsal and superior Anterior Cingulate Cortex, SMA, Dorso-medial PFC, Insula (extracted from the Canlab atlas) and the Putamen, Caudate, Substantia Nigra, Habenula, Globus Pallidus, Red Nuclei (extracted from the Pauli atlas).

**Extra info MRI data acquisition**

MRI data were acquired on a 3 Tesla Philips Achieva scanner, using a 32-channel head coil, at the Department of Radiology of the University Hospitals Leuven. The functional runs (226 volumes each) were recorded using a multi-band sequence (60 axial slices; FOV = 224 x 224 mm; in-plane resolution = 2 x2 mm; interslice gap = 0.2 mm; TR = 2000 ms; TE = 30 ms; MB = 2; flip angle = 90°). In addition, a high resolution T1-weighted anatomical image was acquired for each subject for co-registration and normalization of the EPI data (182 axial slices; FOV = 256 x 240 mm; in-plane resolution = 1 x 1 mm; TR = shortest, TE = 4.6 ms, flip angle = 8°).

**Results from the fasting/re-feeding procedure**

Physical hunger

A 2 (Before-After breakfast) by 2 (Group: Fasting and Re-feeding Group) repeated-measure analysis of variance ANOVA revealed a significant main effect of Group ($F_{(1,47)} = 39.470$, $p < 0.001$, Partial $Eta^2 = .451$), Time ($F_{(1,47)} = 40.288$, $p < 0.001$, Partial $Eta^2 = 0.456$), and Glucose by Group interaction ($F_{(1,47)} = 27.355$, $p < 0.001$, Partial $Eta^2 = 0.363$), with the Re-feeding Group reporting lower levels of physical hunger after breakfast than the Fasting Group ($p < 0.001$).

Emotional hunger

A 2 (Before-After breakfast) by 2 (Group: Fasting and Re-feeding Group) repeated-measure analysis of variance ANOVA revealed a significant main effect of Group ($F_{(1,47)} = 30.323$, $p < 0.001$, Partial $Eta^2 = 0.387$), Time ($F_{(1,47)} = 56.903$, $p < .001$, Partial $Eta^2 = 0.542$), and Glucose by Group interaction ($F_{(1,47)} =79.111$, $p < 0.001$, Partial $Eta^2 = 0.622$), with the Re-feeding Group reporting lower levels of emotional hunger after breakfast than the Fasting Group ($p < 0.001$).

Glucose

A 2 (Before-After breakfast) by 2 (Group: Fasting and Re-feeding Group) repeated-measure analysis of variance ANOVA revealed a significant main effect of Group ($F_{(1,47)} = 28.520$, $p < 0.001$, Partial $Eta^2 = .373$), Time ($F_{(1,47)} = 70.686$, $p < 0.001$, Partial $Eta^2 = 0.596$), and Glucose by Group interaction ($F_{(1,47)} = 96.599$, $p < 0.001$, Partial $Eta^2 = 0.668$), with the Re-feeding Group showing higher levels in glucose after breakfast than the Fasting Group ($p < 0.001$).

**Homogeneity of the two groups**

The participants from the two groups did not differ in terms of age, Body Mass Index (BMI), US intensity selected before the start of the ART, and tolerance to distress as measured by the Distress Tolerance Scale and eating disorders (all p-values > 0.05).

### PAVLOVIAN LEARNING PHASE

**Retrospective US expectancy** The results from the Group (Fasting/Re-feeding) by CS (CS-/CS+$_{av}$/CS+$_{unav}$) by Time (First/Last) LMM analysis showed the expected decrease in the retrospective US expectancy ratings for the CS- compared to the CS+$_{av}$ ($\beta$ = - 6.800, ± 0.759, df = 191.998, t = - 8.954, p < 0.001) and compared to the CS+$_{unav}$ ($\beta$ = - 5.880, ± 0.759, df = 192.000, t = - 7.743, p < 0.001) across the two groups. No significant group-related effects were found, see **figure S1**.

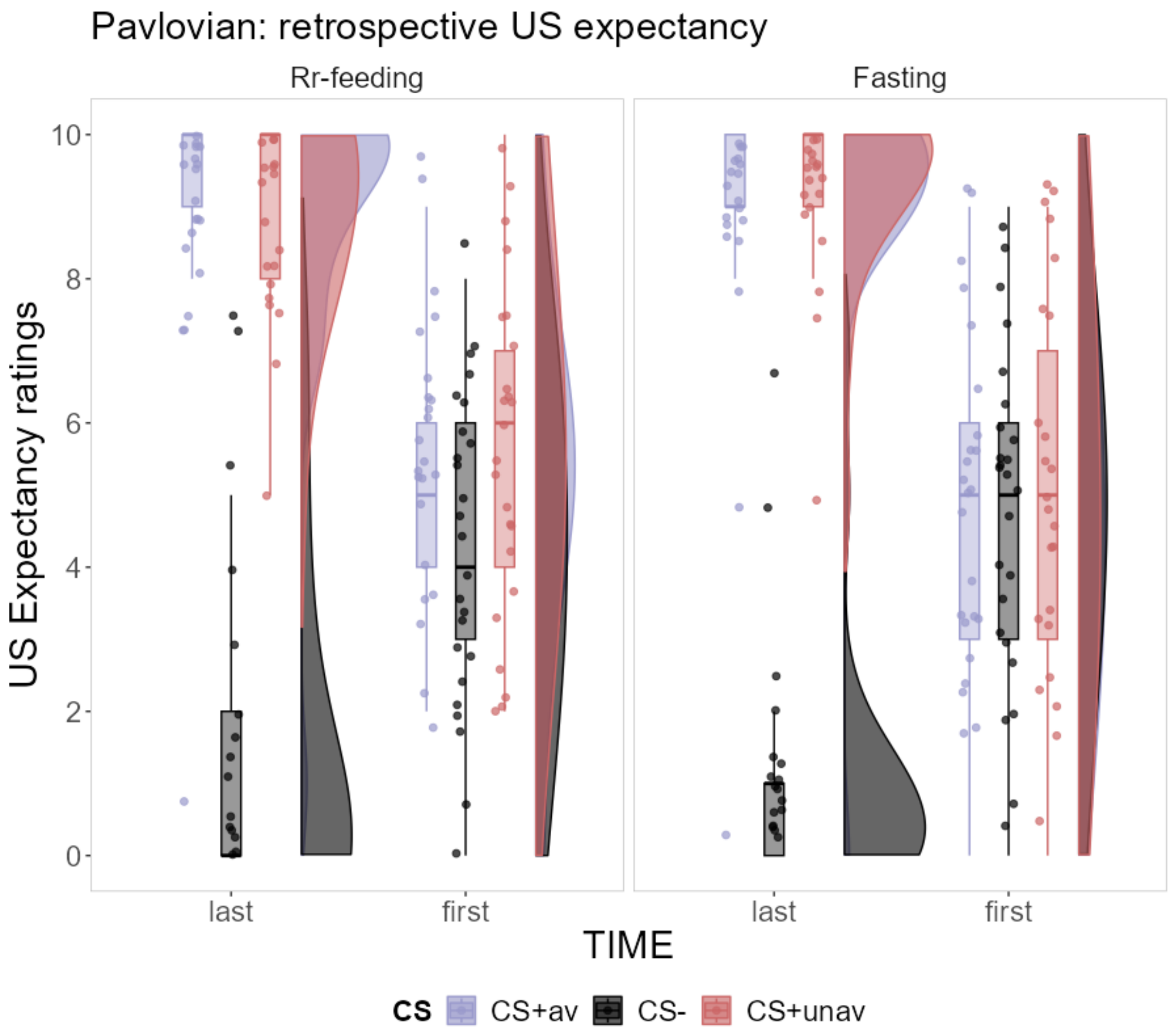


**Figure S1** The retrospective US expectancy ratings collected at the end of the Pavlovian learning phase.

#### Brain activations during US anticipation (extended mask)

**Table S1** The significant brain activations during US anticipation with an initial uncorrected p = 0.001 selected at cluster-level, and a subsequent extent-threshold of a (Family Wise Error, FWE)-corrected p = 0.05.

| Contrast | Name cluster (Brainnetome/MNI Glasser) | Size (cluster) | Peak MNI (x, y, z) | Alpha |
|---|---|---|---|---|
| CS+$_{unav}$ versus CS- | Right Middle Cingulate Gyrus | 420 | -1.5, -11.5, 40.3 | << .01 |

| | | | | |
|---|---|---|---|---|
| | Left Amygdala | 111 | 20.5, 0.5, -14.7 | << .01 |
| | Right Amygdala | 43 | -23.5, -3.5, -10.3 | < .01 |
| | Left Putamen | 28 | 22.5, -5.5, 0.7 | < .02 |
| CS$_{+av}$ versus CS- | Left Middle Cingulate Cortex | 133 | 4.5, -9.5, 40.3 | << .01 |
| | Right Anterior 24 | 128 | - 3.5, -11.5, 42.5 | << .01 |
| | Left Amigdala | 80 | 20.5, 2.5, -14.7 | << .01 |
| | Right Putamen | 26 | -33.5, -3.5, -5.9 | < .01 |
| | Left Putamen | 14 | -32.5, -10.5, -5.9 | < .04 |
| | Right Middle Cingulate Gyrus | 14 | -3.5, -27.5, 31.5 | < .04 |

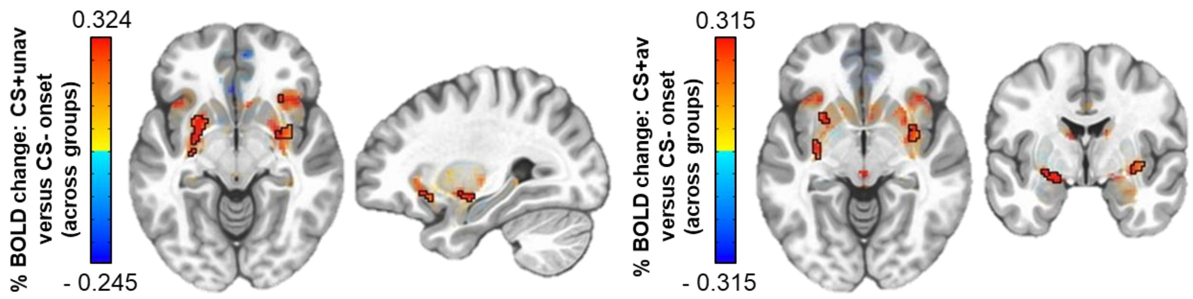


**Figure S2** Brain response to US anticipation (CS$_{+av}$ versus CS- and CS$_{+unav}$ versus CS-) during the Pavlovian learning phase.

**Skin Conductance Responses** For the Pavlovian learning phase we observed a normal distribution of the data. For this reason, we applied an LMM analysis. Since the organization of the presentation of the CSs was set in two blocks, the analysis on SCR to US anticipation was run separately for block 1 and block 2. Given that each block contains only four trials, we averaged the SCR for each type of CS (CS- and CS+) within each block. The results from this analysis showed that during the presentation of the CSs- there was a significantly lower SCR compared to the CSs+ in the first block ($\beta$ = - 0.286, ± 0.072, df = 310.406, t = - 3.947, p < 0.001) as well as in the second block ($\beta$ = - 0.256, ± 0.052, df = 316.066, t = - 4.892, p < 0.001). Importantly, no group differences were found, (all p-values > 0.05).

### AVOIDANCE PHASE

**Retrospective US expectancy** The results from the Group (Fasting/Re-feeding) by CS (CS-/CS$+_{av}$/CS$+_{unav}$) by button press (yes/no) LMM analysis showed the expected lower retrospective US expectancy ratings for the CS$_{+av}$ compared to the CS$_{+unav}$ when the correct avoidance action was made ($\beta$ = - 6.920, ± 0.759, df = 282.000, t = 9.121, p < 0.001), and the expected increase in the retrospective US expectancy ratings for the CS$+_{av}$ when the participant did not press the button (vs did press) (CS by button press: $\beta$ = 5.960, ± 0.759, df = 282.000, t = 7.856, p < 0.001). No significant Group-related effects were found (all p-values > 0.05), see **figure S3**.

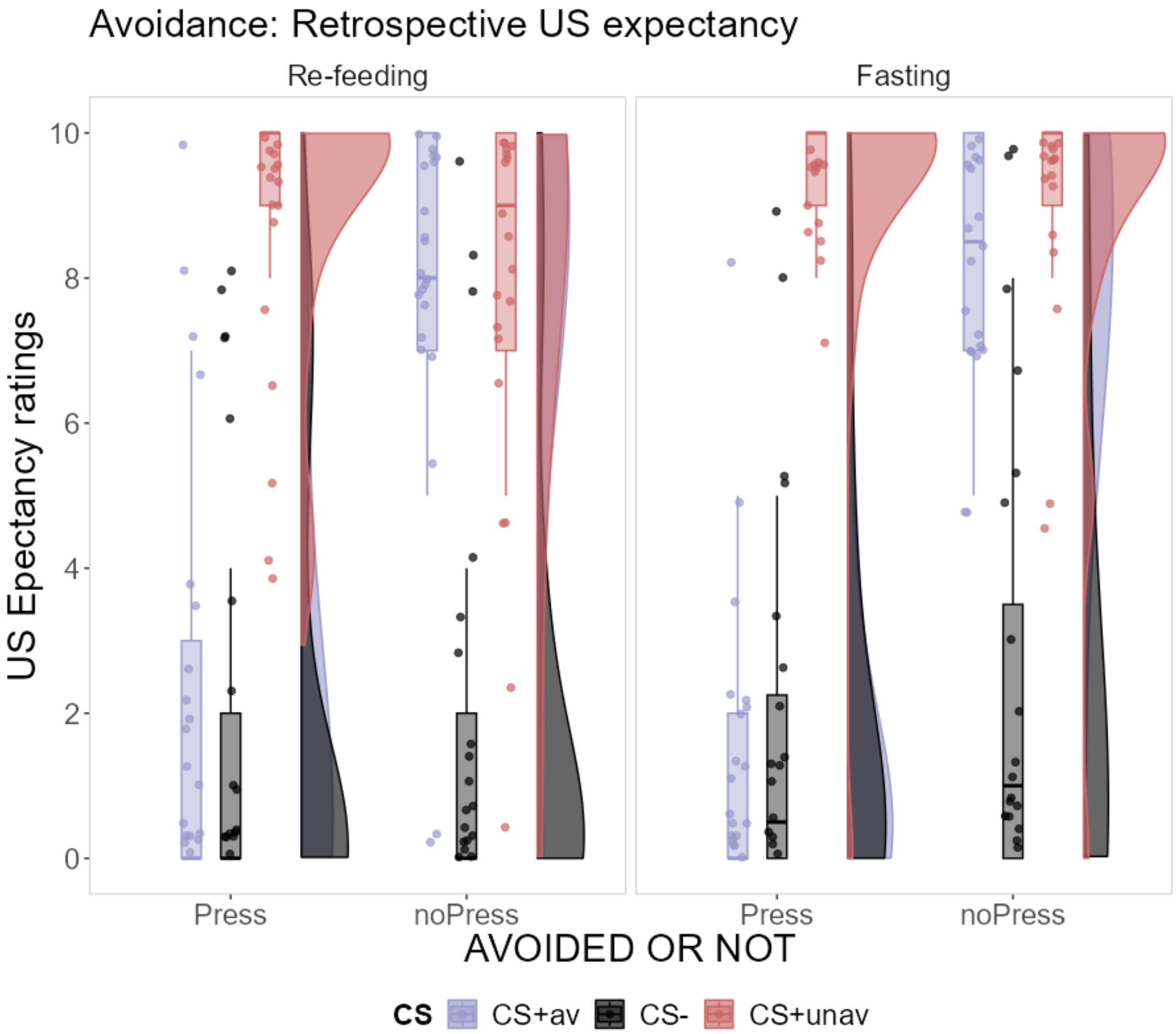


**Figure S3** The retrospective US expectancy ratings collected at the end of the avoidance learning phase.

**ROIs analysis during CS anticipation: no group differences**

There were no significant group differences in the contrast CS- vs $CS_{+av}$ (onset) in Nac, VTA, and vmPFC. The results from the separated independent t-tests showed indeed the absence of any group difference in differential activation in Nac ($p = 0.800$), VTA ($p = 0.248$), and vmPFC ($p = 0.835$).

**Brain activations during US anticipation (extended mask): overall sample**

The presentation of the $CS_{+unav}$ compared to the CS- (onset) elicited activations in the expected brain areas, including the bilateral insula in their anterior parts, and dorsal ACC, see **table S2**, **figure S4**. We did not find significant differences in brain activations in response to the anticipatory phase of the $CS_{+av}$ vs CS-. This result was probably given by the fact that once the participant pressed the button during the $CS_{+av}$, the anticipation of the US omission was no longer able to elicit anticipatory fear and the $CS_{+av}$ was perceived similarly to the safe CS-. As for the other analyses, we used an initial uncorrected $p = 0.001$ and an extent-threshold of a (FWE)-corrected $p = 0.05$).

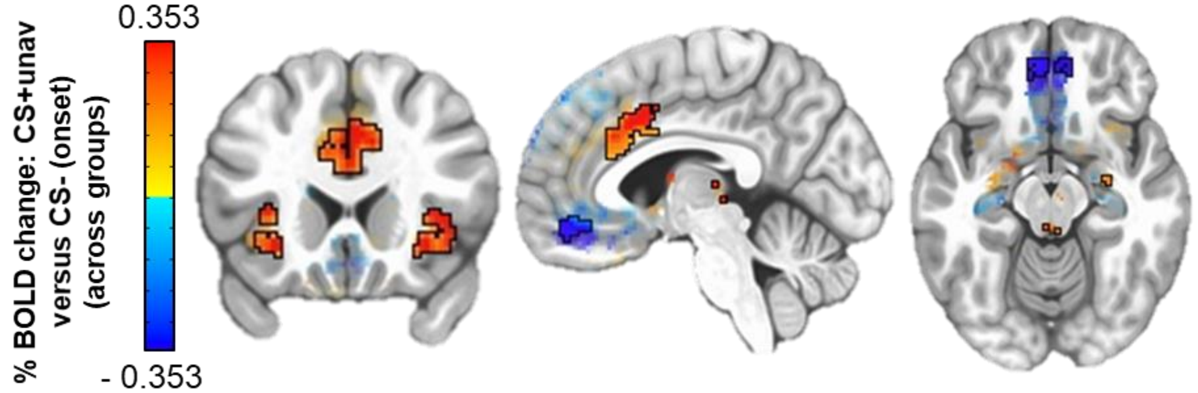


**Figure S4** The brain response to $CS_{+unav}$ versus the CS- (onset) across the two groups during the avoidance learning phase.

**Table S2.** The significant de/activations during the $CS_{+unav}$ versus the CS- (CSs onset) found by the voxel-based analysis during the avoidance learning phase, (initial uncorrected p = 0.001 and an extent-threshold of a (FWE)-corrected p = 0.05).

| **Contrast** | **Name cluster (Brainnetome/MNI Glasser)** | **Size (cluster)** | **Peak MNI (x, y, z)** | **Alpha** |
|---|---|---|---|---|
| $CS_{+unav}$ versus CS- | Right Anterior 24 | 615 | -1.5, -13.5, 40.3 | << .01 |
| | Right anterior Insula | 144 | -33.5, -27.5, 0.7 | << .01 |
| | Left anterior Insula | 86 | 32.5, -21.5, 7.3 | << .01 |
| | Left Orbitofrontal Cortex 10r (negative cluster) | 81 | 2.5, -45.5, -16.9 | << .01 |
| | Right Orbitofrontal Cortex | 38 | -1.5, -45.5, -19.1 | < .01 |
| | Left Thalamus | 28 | 0.5, 32.5, -3.7 | < .02 |
| | Right Caudate | 25 | -11.5, 0.5, 13.9 | < .02 |
| | Left Caudate | 24 | 10.5, -1.5, 9.5 | < .02 |
| | Rigth A32p | 19 | -3.5, -35.5, 24.9 | < .03 |

**Further explorative ROIs analysis during CS offset (overall sample)**

The results from the two independent sample t-tests showed the absence of a significant $CS_{+av}$ versus CS- brain activation in VTA, Nac, and vmPFC (all p-values > 0.05), in line with the results from the analysis reported in the main text.

*VTA* The results from an explorative independent t-test combining the two types of omissions (CS- plus $CS+_{av}$ vs baseline) showed the absence of a significant activation in the VTA, p > 0 .050).

*Nac* The results from an explorative independent t-test combining the two types of omissions (CS- plus $CS+_{av}$ vs baseline) showed a significant activation in the Nac (t = 2.136, df = 43, p-*uncorrected* = 0.038).

*vmPFC* The results from an explorative independent t-test combining the two types of omissi ons (CS- plus $CS+_{av}$ vs baseline) showed a significant activation in the vmPFC (t = 4.088, df = 43, p-*corrected* < 0.001).

**Brain activations during US omissions (extended mask): overall sample**

The presentation of the $CS_{+av}$ compared to baseline and the presentation of the CS- compared to baseline are reported below, **table S3**.

**Table S3** The results from the t-tests in the whole sample for the contrast $CS+_{av}$ vs baseline and CS- vs baseline, run separately for each CS (CS offset) with an initial uncorrected p = 0.001, and a subsequent extent-threshold of an (FWE)-corrected p = 0.05.

| Contrast | Name cluster (Brainnetome/MNI Glasser) | Size (cluster) | Peak MNI (x, y, z) | Alpha |
|---|---|---|---|---|
| $CS+_{av}$ vs baseline | Left SMA | 295 | 2.5, -17.5, 51.3 | << .01 |
| | Left ventral medium Putamen | 125 | 24.5, -3.5, -8.1 | << .01 |
| | Left SMA | 110 | -1.5, -55.5, 13.9 | << .01 |
| | Left Middle Insula | 107 | 40.5, -19.5, -3.7 | << .01 |
| | Right Superior Medial Gyrus (A10) | 107 | -1.5, -13.5, 53.5 | << .01 |
| | Right Superior Frontal Gyrus (A8) | 73 | -21.5, -35.5, 55.7 | << .01 |
| | Left Superior Medial Gyrus (A9) | 52 | -1.5, -31.5, 40.3 | < .01 |
| | Right Putamen/Globus Pallidum | 39 | -23.5, 0.5, 7.3 | < .01 |
| | Right Anterior ventral Insula | 27 | -31.5, -25.5, - 1.5 | < .02 |
| | Right ACC (A32sg) | 24 | -11.5, -43.5, 9.5 | < .02 |
| | Left Superior Medial Gyrus (A9) | 23 | 6.5, -61.5, 33.7 | < .02 |
| | Right Superior Frontal Gyrus (A9) | 23 | -15.5, -47.5, 46.9 | < .02 |
| | Right Middle Insula | 17 | -39.5, -19.5, 0.7 | < 0.4 |
| CS- vs baseline | Left Putamen/Globus Pallidum | 149 | 20.5, -7.5, 2.9 | << .01 |
| | Left SMA (A8) | 121 | -3.5, -11.5, 44.7 | << .01 |
| | Right ventral medial Putamen | 93 | -23.5, -7.5, 2.9 | << .01 |
| | Left ACC (A24, Ap32) | 77 | 10.5, -19.5, 29.3 | << .01 |
| | Left SMA(A8) | 54 | -1.5, -13.5, 53.5 | < .01 |
| | Right Caudate Nucleus | 42 | -11.5, 0.5, 16.1 | < .01 |
| | Left Caudate Nucleus | 38 | 10.5, -5.5, 16.1 | < .01 |
| | Left SMA | 37 | 2.5, -19.5, 51.3 | < .01 |
| | Left dorsal lateral Putamen | 36 | 28.5, 2.5, -8.1 | < .01 |
| | Left Superior Medial Gyrus | 28 | 2.5, -31.5, 40.3 | < .01 |
| | Right Insula | 22 | -31.5, -15.5, 7.3 | < .02 |
| | Left Caudate | 16 | 14.5, -13.5, 7.3 | < .04 |
| | Left Superior Medial Gyrus (A9) | 15 | 2.5, -57.5, 16.1 | < .05 |

**Skin Conductance Response during US anticipation**

The results from the Group (FG/RG) by CS($CS-/CS+_{av}$) by Trial GLMM analysis (logit component) did not show any significant group effect.

In line with fear extinction learning paradigms, we found the expected significant reduction in the probability to respond during the CS- compared to the $CS_{+av}$ ($\beta$ = - 0.635, ± 0.228, - 2.780, p = 0.005), and during the CS- compared to the $CS_{+unav}$ ($p < 0.001$), as well as a significant general reduction in the probability of SC responding over the course of the learning phase (Trial: $\beta$ = - 0.200, ± 0.046, z = - 4.341, $p < 0.001$).

### EXTINCTION PHASE

**ROIs analysis during CS anticipation: no group differences**

We did not find any significant group difference for the CS- vs $CS_{+av}$ (onset) contrast in Nac, VTA, and vmPFC. The results from the separated independent t-tests showed indeed the absence of any group difference in this differential activation in Nac (p = 0.833), VTA (p = 0.248), and vmPFC (p = 0.879).

**Brain activations during US anticipation (extended mask): overall sample**

We did not find any significant difference in brain activity when we compared the $CS_{+av}$ versus the CS- onset. Also, during anticipation of the US, similarly to the US omission, the $CS_{+av}$ and CS- presentation induced similar brain activity (probably given by the change in context, see the 'Limits' section in the main text). This included a strong deactivation of the vmPFC, Nac bilaterally, and positive activation of the anterior ventral Insula, Putamen bilaterally, and dorsal ACC. These activations were found using an initial uncorrected p = 0.001 and an extent-threshold with an (FWE)-corrected p = 0.05), **figure S5**.

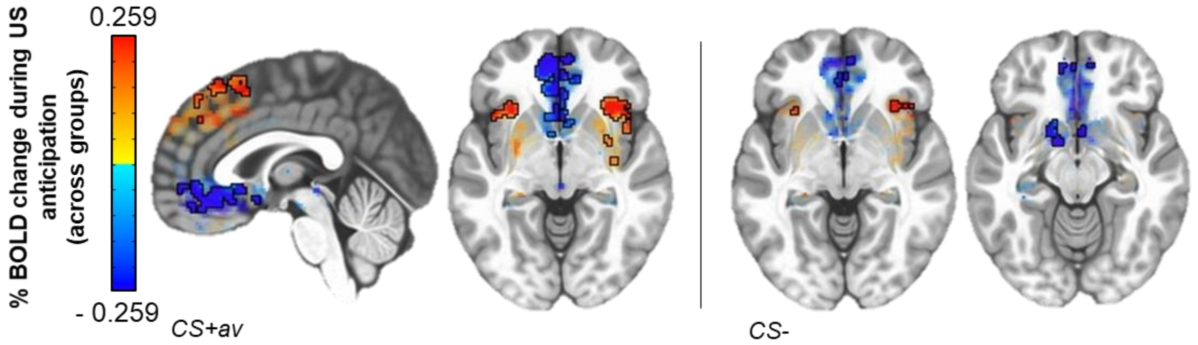


**Figure S5** Brain de/ activations during US anticipation for each CS (vs baseline) across the two groups during the fear extinction paradigm.

**Table S4** The significant de/activations during the CS+av as well as for the CS- vs baseline during the anticipatory phase of the fear extinction learning paradigm. We used an initial uncorrected p = 0.001, and a subsequent extent-threshold of an (FWE)-corrected p = 0.05.

| **Contrast** | **Name cluster (Brainnetome/MNI Glasser)** | **Size (cluster)** | **Peak MNI (x, y, z)** | **Alpha** |
|---|---|---|---|---|
| CS+av vs baseline | Left 10d Mid Orbital Cortex (negative cluster) | 343 | 2.5, - 53.5, - 3.7 | << .01 |
| | Right Insula | 227 | - 43.5, - 19.5, - 1.5 | << .01 |
| | Right p32 Mid Orbital Cortex | 181 | - 1.5, - 53.5, - 5.9 | << .01 |
| | Left Putamen | 81 | 24.5, - 1.5, 5.1 | << .01 |
| | Left Middle CIngulate Cortex | 69 | 8.5, - 7.5, 40.3 | << .01 |
| | Left Insula | 65 | 30.5, - 21.5, - 8.1 | << .01 |
| | Right SMA | 63 | - 1.5, - 15.5, 53.5 | << .01 |
| | Left SMA | 41 | - 1.5, - 29.5, 57.9 | < .01 |
| | Left SMA | 17 | - 3.5, - 15.5, 40.3 | < .04 |
| | Right Putamen | 15 | - 21.5, - 1.5, 7.3 | <.05 |
| CS- vs baseline | Left Mid Orbital Gyrus | 56 | 8.5, - 49.5, - 8.1 | < .01 |
| | Right Insula | 38 | - 41.5, - 21.5, - 3.7 | < .01 |
| | Left Piriform Cortex/amygdala (negative cluster) | 36 | 16.5, - 3.5, - 14.7 | < .01 |
| | Left SMA | 27 | 4.5, - 9.5, 42.5 | < .02 |

**Further explorative ROIs analysis during CS offset: overall sample**

The results from the two independent sample t-tests showed the absence of a significant $CS_{+av}$ versus CS- brain activation in VTA, Nac, and vmPFC (all p-values > 0.05), in line with the results from the analysis reported in the main text.

*VTA* The results from an explorative independent t-test combining the two types of omissions (CS- plus $CS+_{av}$ vs baseline) showed a significant activation in the VTA ($t = 3.2447$, $df = 42$, p-*corrected* = 0.002).

*Nac* Similarly to the avoidance learning phase the results from an explorative independent t-t est combining the two types of omissions (CS- plus $CS+_{av}$ vs baseline) showed a significant activation in the Nac ($t = 2.6118$, $df = 42$, p-*corrected* = 0.012).

*vmPFC* Differently from the VTA and Nac, no significant results were found when using the $CS_{+av}$ plus CS- versus baseline contrast in one independent t-test for this ROI ($t = 1.305$, $df = 45$, $p = 0.099$). Although this contrast is not specific for the type of US omission (CS-specific) we expected the vmPFC to be active during a general period of safety. From a close visual inspection of the results from the extended mask, we, however, observed the presence of two sub/clusters of opposite valence which might have canceled out part of the overall activation when using the averaged beta weights for the t-test, see **figure S6**. To further characterize the behavior of this region during general US omissions we run an explorative voxel-wise analysis of this large ROI in AFNI. As we expected, the results indicated that the mask included a significant positive cluster in VmPFC when using the combined contrast CS- plus $CS_{+av}$ versus the baseline (133 voxels, peak x: 2.5, y: -53.5, z: 5.1); and a negative but no significant sub-cluster in the most orbital part of the vmPFC.

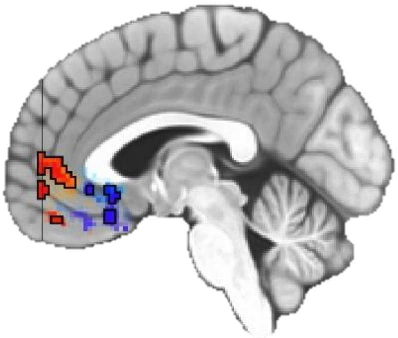

**Figure S6** The opposite clusters found in the vmPFC mask.

**Table S5** The results from the independent t-tests across groups for the contrast $CS_{+av}$ vs baseline and CS- vs baseline run separately across the whole sample during the US omissions of the fear extinction learning paradigm. Extended corrected threshold alpha below (FWE-corr) .05 with an initial $p = 0.001$ uncorrected).

| Contrast | Name cluster (Brainnetome/MNI Glasser) | Size (cluster) | Peak MNI (x, y, z) | Alpha |
|---|---|---|---|---|
| $CS+_{av}$ vs baseline | Left SMA | 845 | -1.5, -15.5, 42.5 | << .01 |
| | Left Putamen | 109 | 24.5, -7.5, 2.9 | << .01 |
| | Right Superior Frontal Gyrus (A9l) | 77 | -15.5, -43.5, 51.3 | << .01 |
| | Right Caudate | 48 | -13.5, -13.5, 5.1 | << .01 |
| | Right Putamen | 42 | -23.5, -7.5, 5.1 | << .01 |
| | Left Superior Medial Gyrus (A10m) | 39 | -1.5, -55.5, 20.5 | << .01 |
| | Left Caudate | 31 | 16.5, -17.5, 5.1 | < .01 |

| | | | | |
|---|---|---|---|---|
| | Left Middle Insula | 21 | 32.5, -19,5, 7.3 | < .02 |
| | Left Superior Medial Gyrus (A10m) | 21 | 4.5, -59.5, 29.3 | < .02 |
| | Right Middle Insula | 20 | -35.5, -17.5, 5.1 | < .02 |
| | Left ACC (A32p) | 16 | 8.5, -43.5, 18.3 | < .04 |
| | Left Superior Medial Gyrus (A10m) | 15 | 4.5, -57.5, 9.5 | < .04 |
| | Left Superior Frontal Gyrus | 13 | 24.5, -33.5, 55.7 | < .05 |
| CS- vs baseline | Left SMA (A8) | 220 | 8.5, -9.5, 44.7 | << .01 |
| | Left Putamen | 147 | 24.5, .5, 9.5 | << .01 |
| | Left SMA (A8) | 146 | -1.5, -13.5, 44.7 | << .01 |
| | Right Insula | 21 | -31.5, -15.5, 7.3 | < .02 |
| | Right ACC (A32p) | 17 | -9.5, -29.5, 24.9 | < .03 |
| | Left Superior Medial Gyrus (A9m) | 16 | 2.5, -49.5, 44.7 | < .04 |
| | Right Hyppocampus (CA2) | 13 | -31.5, 14.5, -16.9 | < .05 |

**Retrospective US expectancy**

The results from the Group (FG/RG) by CS (CS-/CS+$_{av}$/CS+$_{unav}$) by Trial (First/Last) LMM analysis showed the expected reduction in the retrospective US expectancy ratings across the CSs (Time: p = 0.001), see **figure S7**.

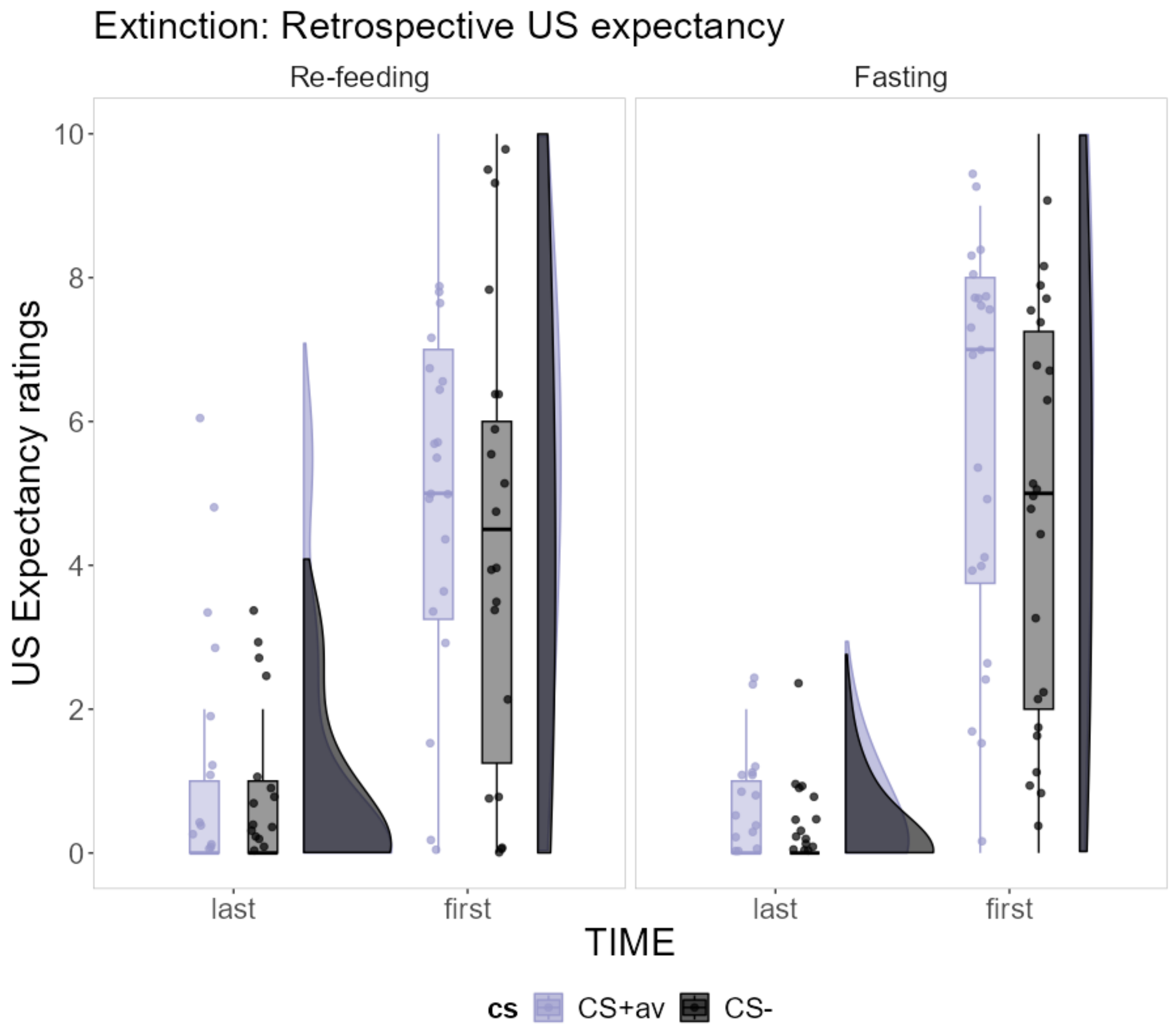

**Figure S7** The retrospective US expectancy ratings collected at the end of the fear extinction learning phase.

#### Skin Conductance Responses during US anticipation

The results from the Group (FG/RG) by CS(CS-/CS+$_{av}$) by Trial GLMM analysis did not show any significant group related effect. In general, we observed the expected significant reduction in SCRs over the presentation of the CSs (Trial: $\beta$ = - 0.331, ± 0.086, z = - 3.828, p < 0.001).

#### Neuropsychological questionnaires

A 2 (Time: Pre[screening day 1] and Post[post MRI]) by 2 (Group: Fasting and Re-feeding Group) repeated-measure analysis of variance ANOVA revealed the absence of any significant main effect of Group or Group by Time interaction in the Digit Span, DS (Forward [MAX: maximum sequence remembered, and CORR: number of correct answers] and Backward [MAX and CORR]), Need for Cognition Scale, and Psycho Visual Task (all p-values > 0.05). This result indicates that the group differences indicated in the main text were not secondary to any cognitive change following fasting. We only found a Time effect for the maximum sequence remembered for the DS backward (p = 0.016) which might have depend on training-effects on working memory. This effect was however not related to Group.
